# DNA-templated Chiral Metamaterial Array as Information Bits

**DOI:** 10.64898/2026.01.25.701623

**Authors:** Deeksha Satyabola, Shuai Feng, Prathamesh Chopade, Abhay Prasad, Lihui Wang, Nimarpreet Bamrah, Liangxiao Chen, Subhajit Roy, Aaron Sakai, Chao Wang, Petr Sulc, Rizal Hariadi, Hao Yan, Sui Yang

## Abstract

The growth of digital information demands physically secure information bits that combine intrinsic randomness with multi-layered optical readout. Digital metamaterials with physical unclonable function characteristics are promising, but nearly all examples operate at microwave or terahertz frequencies. Extending digital metamaterials to the visible range requires a physical property that allows binary (0/1) encoding via nanoscale geometry and is orthogonal to intensity-or color-based imaging. Chiral plasmonic metamaterials satisfy these criteria perfectly: their broken mirror symmetry yields chirality polarity in the visible spectrum whose sign and magnitude directly encode bit values (“0”, spin-up; “1”, spin-down), while remaining invisible under linear polarization. Here, we realize visible-range digital metamaterials by programming self-assembled DNA origami templates with asymmetrically placed gold nanorods to create discrete 3D chiral metamolecules. Using on-surface DNA origami assembly and the Silica Microsphere-Assisted Patterning by Liquid Elimination (SiMPLE) method, we fabricate large-scale bit arrays on optical active glass substrate in solution with ∼1.33 μm lattice spacing, ∼86% site occupancy, and ∼12-month stability after silicification. These bottom-up fabricated digital metamaterial array exhibits two independent security layers: (1) a macroscopic spatial pattern only visible by dark-field microscopy, and (2) hidden binary information stored in the single-particle chiroptical response, read out by position-resolved circular dichroism spectroscopy. Quantitative analysis confirms reliable bit encoding and high optical randomness arising from slight structural variations. By leveraging the polarity of plasmonic chirality to translate molecular-scale handedness into robust visible-range digital signals, this work establishes a scalable nanophotonic platform for secure optical information storage, authentication, and encryption.

## Introduction

Information bits are basic binary states (0 or 1) encoded in the physical state of a nanoscale object^1–6^, and they are central to next generation optical information storage^7,8^, authentication^9,10^, and physical unclonable devices^11–13^. Since their theoretical introduction in 2014, digital metamaterials^14^ have offered a way to realize such bits, in which each meta-molecule serves as an information bit by having its electromagnetic response discretized into two distinct states. Currently, most experimentally realized digital metamaterials operate within microwave^15,16^, terahertz^17–19^ and acoustic frequencies^20,21^, where binary states are typically encoded via differences in reflection phase or transmission amplitude^22–26^. Extending this concept to the visible optical range has remained elusive^27^, primarily because achieving switchable and single nanoparticle readable optical contrast at visible wavelengths demands sub-10 nm^28,29^ control over three-dimensional (3D) geometry in objects.

Chiral plasmonic metamaterials offer an ideal solution. By breaking mirror symmetry^30,31^, 3D arrangements of plasmonic nanoparticles generate chiral optical response in the visible spectrum^30–35^, with sign and magnitude directly determined by nanoscale handedness. Critically, left- and right-handed configurations of otherwise identical nanoparticles produce opposite circular dichroism (CD) responses^35–40^ while remaining indistinguishable under linear polarization or intensity-based imaging. This intrinsic orthogonality enables robust binary encoding (“0”, spin-up; “1”, spin-down) that is hidden from conventional microscopy yet accessible via single-particle CD spectroscopy. Moreover, slight stochastic variations in nanoparticle angle or position during assembly introduce uncontrollable optical randomness^41,42^, perfect for generating random unclonable labels.

DNA origami^43^ provides the ultimate tool for implementing such chiral digital metamaterials. Its addressable, sub-nanometer precision allows programmable placement of gold nanoparticles into arbitrary 3D chiral geometries in solution^44–46^. The organization of DNA helices in square or honeycomb lattices allows center-to-center spacing of ∼ 2–2.5 nm, greater design flexibility than top-down lithography, for rapid prototyping of complex nanoarchitectures^47,48^. Although electron-beam lithography and extreme ultraviolet lithography remain indispensable for wafer-scale semiconductor manufacturing, they are ill-suited for low-volume, high-complexity 3D plasmonic prototyping where molecular-level structural control is paramount^49,50^. DNA-based self-assembly, by contrast, excels exactly in this regime, complementing with conventional nanofabrication.

Here, using DNA origami self-assembly, we extent the chiral digital metamaterial platform into the visible range. We design DNA origami templates that position gold nanorods into symmetry-broken 3D chiral dimers, each functioning as an independent information bit whose binary state is encoded in the sign of its plasmonic CD signal. For on-surface DNA origami placement, we developed the Silica-Microsphere-Assisted Patterning by Liquid Elimination (SiMPLE) method^51^, using which we organize thousands of these chiral bits into micron-scale arrays with ∼1.33 μm lattice spacing and ∼46% chiral bit occupancy. The resulting arrays were silicified, conferring long-term stability (>12 months) and protection against sustained light and thermal stress during optical measurements, while preserving single nanoparticle chiroptical activity. The resulting platform functions as a two-layer optical authentication system: the first layer reveals a macroscopic spatial pattern via dark-field microscopy, and the second hidden layer stores binary information accessible only through position-resolved CD spectroscopy. By translating molecular-scale handedness into robust, readable digital signals at visible frequencies, this work could guide building of scalable nanophotonic framework for creating random bits for anti-counterfeiting labels^52, 53^, and physical unclonable functions^54–56^.

## Results and Discussion

### Design and Overview of One-bit Chiral Metamaterial Platform

**Figure 1** provides an overview of our chiral digital metamaterial array platform, which integrates bottom-up fabrication strategy for information encoding with single-particle circular dichroism (CD) optical readout for information retrieval. Figure 1A illustrates the synthesis of DNA-origami triangle by self-assembly of a single-strand M13 scaffold with several hundreds of short-DNA strands called staples. These DNA-origami triangles contain rationally designed complementary handle sequences that enable subsequent assembly with gold nanorods (AuNRs). The resulting chiral metamaterials are then deposited onto spatially patterned arrays. Owing to the locally symmetry-breaking architecture of chiral metamaterial at each site, differential absorption in left and right circularly polarized light is observed, allowing optical discrimination between left and right chiral metamaterials. Figure 1B illustrates the left chiral metamaterial, which responds to counter-clockwise (spin-up) electric-field polarization and is numerically assigned a 0-bit, and the right-handed chiral metamaterial, which responds to clockwise (spin-down) circular polarization and is assigned a 1-bit. This binary optical response enables the construction of surface-encoded binary information, forming a one-bit digital metamaterial platform. Distinct from conventional top-down digital metamaterials^24^, our platform encodes binary information in the local three-dimensional (3D) orientation of chiral metamaterials^41,42^. A micron-scale separation between neighboring bits prevents ensemble averaging and minimizes interference from adjacent signals. The programmability and modularity of DNA origami for organizing plasmonic nanoparticles, combined with parallel bottom-up pattern fabrication, large-scale organization and compatible integration with optical set-ups, enabled the use of this platform for proof-of-concept random information encoding.

**Figure 1.**
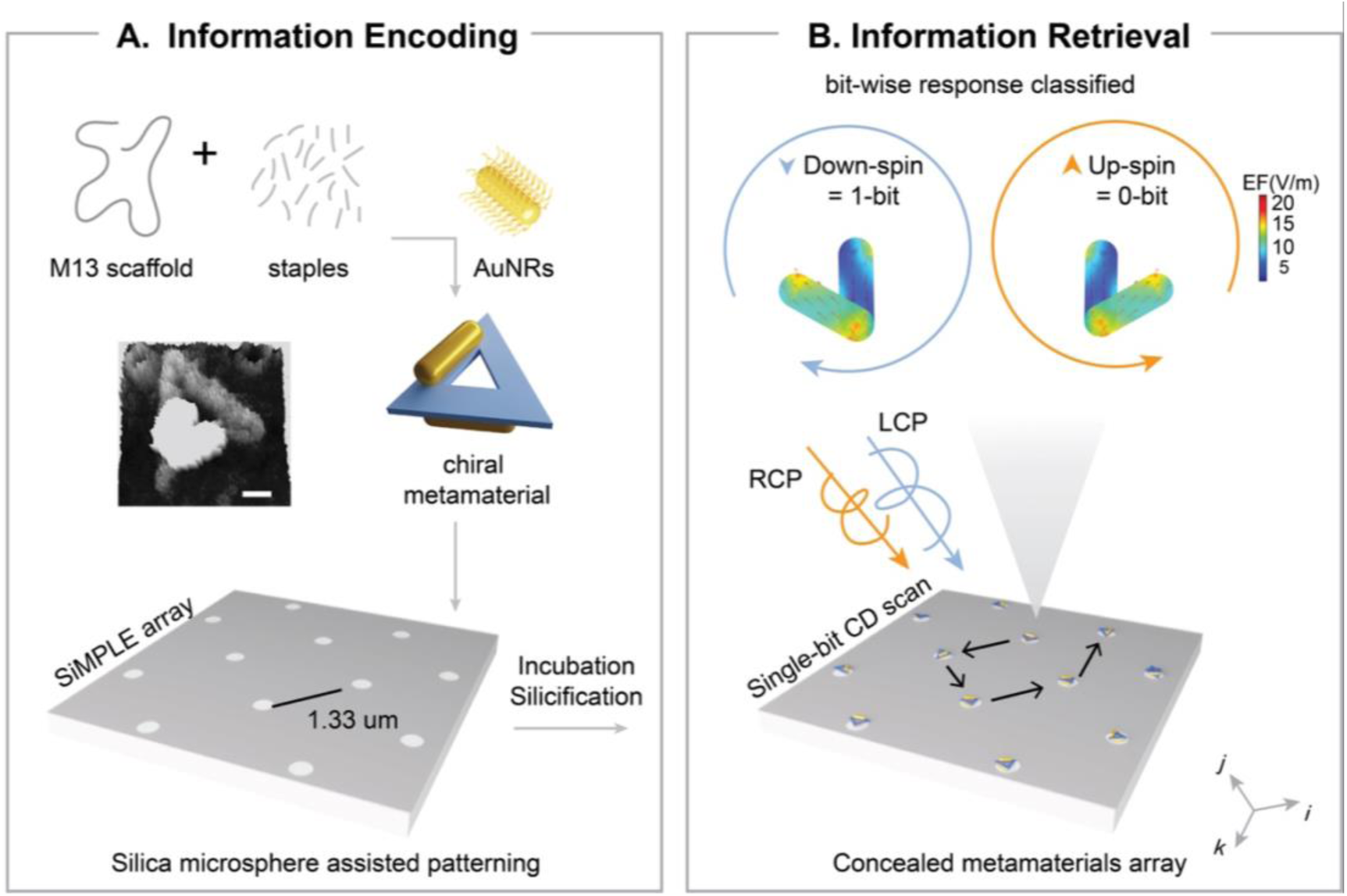
Overview of information encoding and retrieval on chiral coding metamaterial array. A. Synthesis of chiral metamaterials by self-assembly of DNA origami triangle and gold nanorods, followed by random placement on patterned array and silicification; B. Circularly polarized light (LCP, RCP) interacts with the spatially patterned chiral metamaterials bitwise resulting in a distinct CD lineshape (spin), categorizing the bits into 1 or 0.

### Fabrication of Large-scale Single-Molecule Chiral Metamaterial Array

A bottom-up Nanosphere Lithography strategy called Silica Microsphere-assisted Patterning via Liquid Elimination (SiMPLE)^51^ was developed to create a pattered surface (please check SI Section 1, **Figure S1** for detailed procedure). Silica microspheres were chosen due to their higher Young’s Modulus compared to polystyrene nanoparticles used previously in literature. Hertzian contact theory^57^ predicts a directly proportional relationship between pressure applied by nanoparticle on surface (P) and generated spot size (a). We used this relationship and hypothesized that the stiffer and weakly elastic silica microspheres will impart high intermolecular pressure on surface and result in a smaller spot size (Section 1 in Methods).

Atomic force microscopy (AFM) confirmed this hypothesis: for same size distribution of silica and polystyrene nanoparticles (∼1 μm), silica–substrate contact areas were measured as three times smaller than polystyrene (please check SI Section 1 for calculations and **Figure S2**). This indicates that if a micron separated binding spot generated by silica occupies single origami (∼100 nm in diameter), then a binding spot generated by polystyrene will occupy three times more origami nanostructures. Therefore, silica microspheres turned out to be an ideal choice for rapidly generating spatially separated singly occupying array. SiMPLE also expands the capabilities for single-molecule biophysics by resulting in a non-deformative, high-throughput patterning with precisely controlled spot size and well-defined spacing between spots, obliviating issues like stamp deformation faced with techniques like microcontact printing. Furthermore, all steps from DNA assembly, silicification can be performed in liquid state, and thus samples can be processed in batches and scaled in future, allowing for more accessible nanostructure fabrication.

After successfully achieving periodic hydrophilic binding spots on array, we started with deposition of DNA origami triangles on substrate. The 2D origami triangle was observed to be planar with 120 nm edge length in oxDNA simulation (**Figure S3**). A uniform placement of triangle origami was observed facilitated by hydrogen bonding between the negatively charged phosphate backbone of origami and hydroxyl-containing hydrophilic binding spots on substrate in presence of divalent Mg^2+^ ions. A detailed procedure of origami placement is described in SI Section 1. **Figure S4** shows a uniform occupancy of triangle origami on array, which was further validated by high-resolution DNA-PAINT imaging. A concentrations screening of origami was performed using DNA-PANT, each triangle origami contained 12 handles for cyanine 3 (cy3) imager which was clearly visible in the zoomed in image Figure S4. A detailed sample preparation and imaging protocol is described SI Section 1. PAINT images (SI **Figure S5**) revealed a ∼ 45-50% occupancy of single-origami triangles, ∼25% dimer-origami triangles, ∼15% aggregates/multi-origami triangles and ∼10-15% spots without any origami triangles (empty).

Next, largescale patterning was confirmed using photobleaching, where triangle origami with 12 overhangs were hybridized with cy3. Photobleaching analysis predominantly showed single-step photobleaching events (<10/12 steps), also validating single-triangle occupancy on array. (SI **Figure S6**).

After validating large-area single occupancy of DNA origami triangles as a model system, chiral metamaterial arrays were fabricated. As a proof-of-concept, a symmetry breaking conformation of AuNRs on planar DNA-origami triangle was synthesized using low-pH method as described in SI (**Figure S7**). In this conformation, the AuNRs adopt a V-shaped arrangement with approximately 60° inter-rod angle to achieve strongest dipole coupling, as described in previous section. The assembly of chiral metamaterials was confirmed by negative-stain TEM (SI **Figure S7**). These assemblies were incubated on array at different concentrations to optimize single-molecule occupancy, with 120 pM and a 1-hour incubation yielded optimal result. The assembled chiral metamaterial array was then washed with multiple buffer rinses and silicified to stabilize the angle between the nanorods for further optical measurements. Initially, silicification was performed on randomly deposited triangle origami and chiral metamaterials at 1% TEOS/TMAPS on the same optically transparent substrate on which array is prepared (**Figure S8, S9**). After confirming conditions for silicification on glass substrate, similar procedure was followed on array at different concentrations of TEOS/TMAPS such as 0.25, 0.50 and 1 % as shown in **Figure S10**, among these 0.5 and 1% was found obscure the AuNRs angular feature on SEM. Therefore, subsequent studies were performed with 0.25% TEOS/TMAPS for 4 hrs. The silicified chiral metamaterial arrays were then thoroughly washed, ethanol-dried and large-scale occupancy was confirmed by SEM (Figure 2A**–C**).

**Figure 2.**
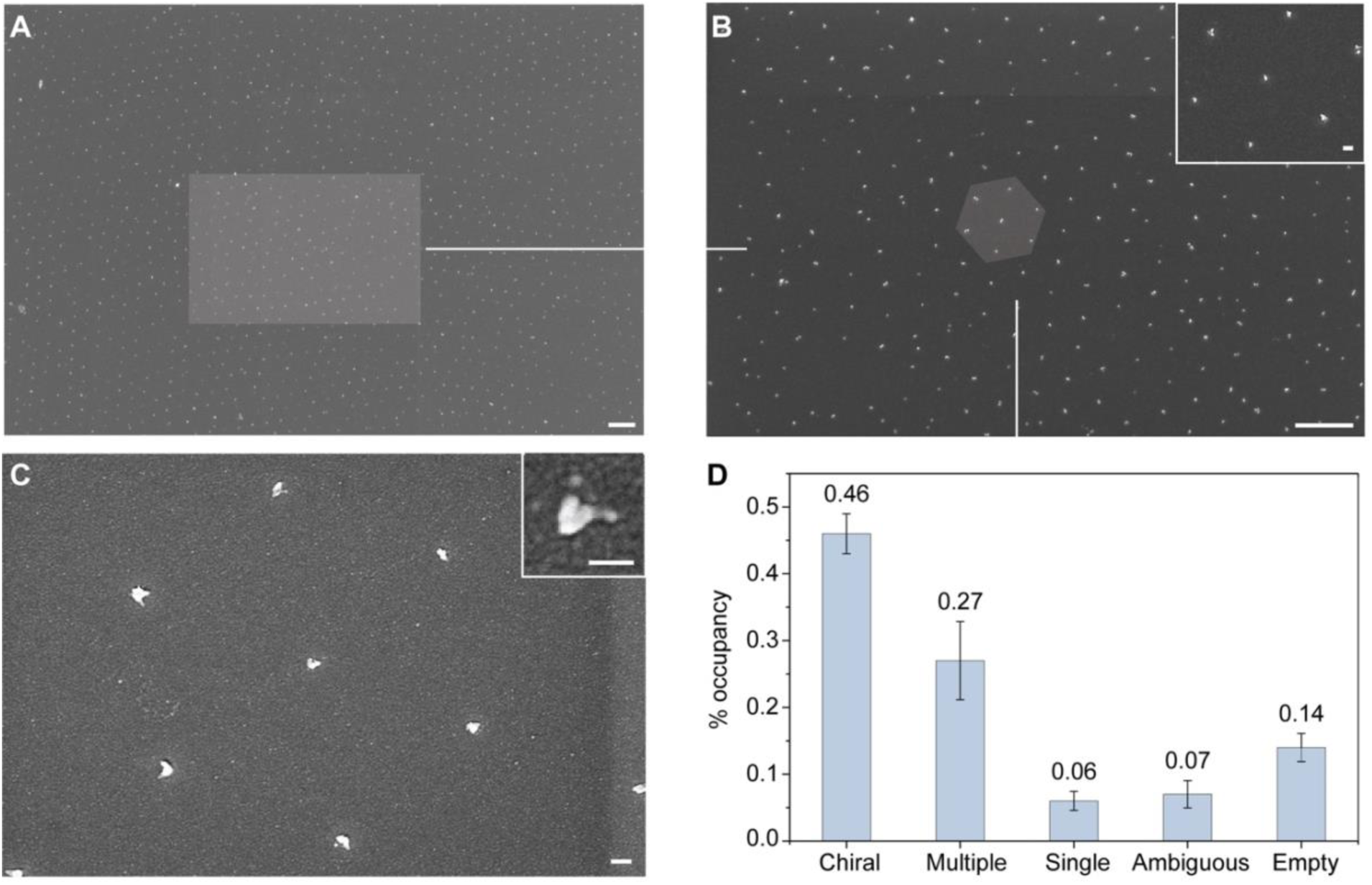
Largescale chiral metamaterial array. (A–C) SEM images showing largescale patterning of chiral metamaterials in a hexagonal pattern. (D) Occupancy of various geometries of chiral metamaterials obtained in SEM after silicification. Scale bar for A–B is 2 𝜇m and C is 100 nm (inset are 100 nm).

In a randomly distributed non-silicified and silicified chiral metamaterials, we counted ∼500 particles and observed an improvement in 60 angle between two AuNRs after silicification (**Figure S11**). The counted particles were categorized as single, dimer/chiral, and multimer AuNRs (Figure 2D). Chiral particles were further classified into < 20 and between >20 and ≤60, which tend to stay in the later form by at least 10% improvement after rigidifying the constructs through silicification (**Figure S11**). Chiral AuNRs showed the strongest anisotropic signal in simulation studies (**Figure SI.** Simulation of single, dimer and multimer AuNRs comparison), demonstrating high fidelity to encode and retrieve chiroptical information.

### Bit-by-bit CD Information Measurements

Correlated single-particle spectroscopy and SEM imaging method optical response was used to directly link optical readout with its corresponding structural geometry (**Figure S12**).

Measurements were performed using a high-angle dark-field microscope with sample sandwiched between the objectives. Detailed instrument set-up is described in Methods Section. Electromagnetic wave frequency domain (EWFD) simulations were conducted to predicted CD lineshapes of representative populations of chiral dimers. To simplify the readout processes, the overall polarity of lineshapes were used to assign bits which are independent of angles between AuNRs dimers. A positive polarity is assigned as 1-bit, while negative response corresponded to 0-bit as confirmed through theoretical study in Figure 3A. The simulation confirm that polarity does not flip or degrade with angle variations. Large angles produce narrower CD responses, while small angles produce broader and more intense ones, but the sign stays ties to the chiral handedness, which make it easier to read bits in mixed or imperfect dimer assemblies without angle-specific calibrations.

**Figure 3.**
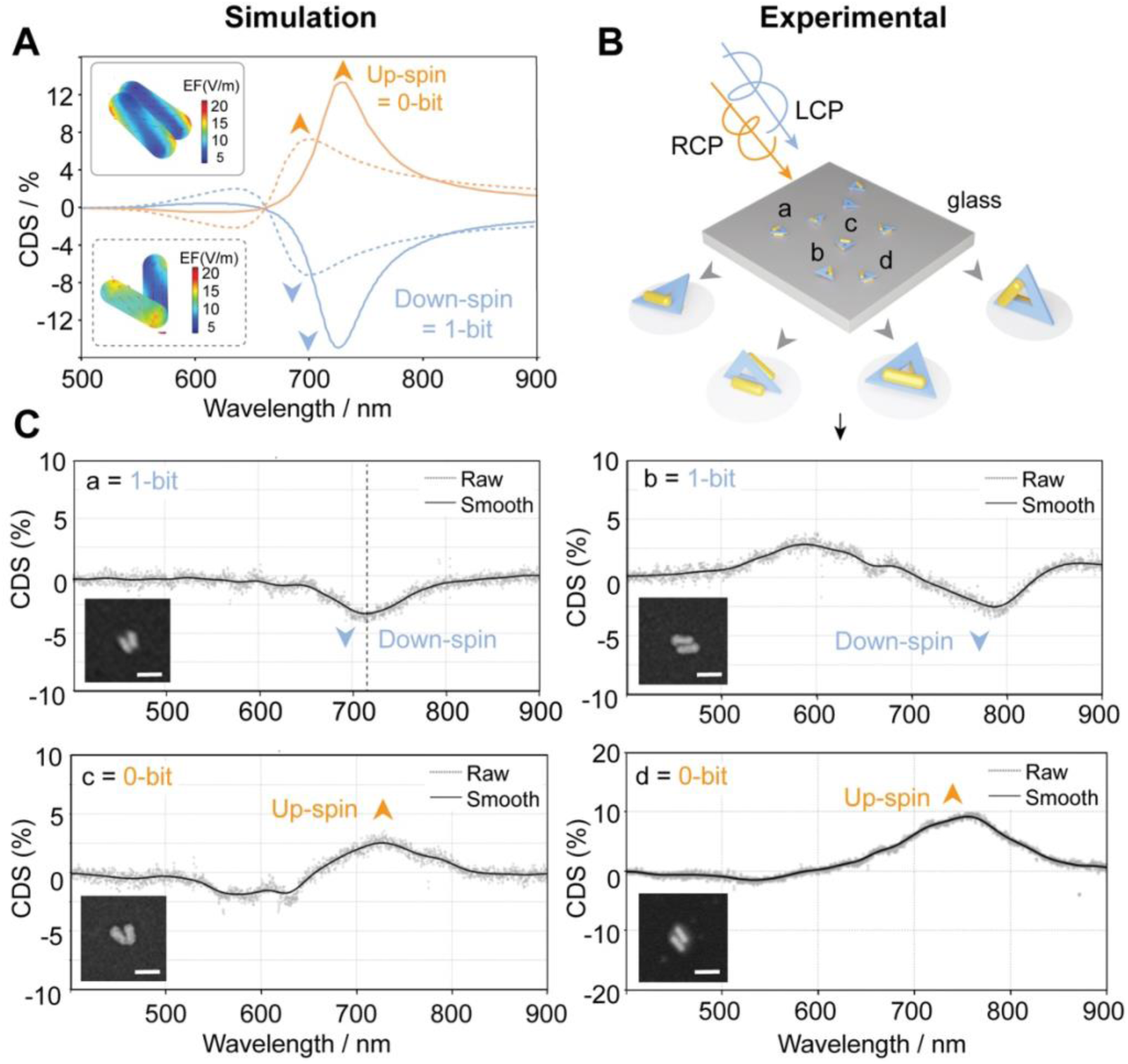
Single-molecule circular dichroism measurements and bit assignment. (A) Electromagnetic wave, frequency domain simulations of two representative dimers showing clockwise up-spin and anti-clockwise down-spin characteristics of chiral metamaterials on transparent glass substrate, Spin is independent of angle between AuNRs and only depend on geometry. (B) Circular dichroism is measured experimentally from chiral metamaterials on substrate. (C) Representative experimental CD lineshapes from single chiral metamaterials on surface defined as 0-bit or 1-bit based on spin. EWFD simulations of the two arrangements correlated well with experimental CDS. Scale bar are 100 nm.

Experimentally, we observed angle-independent chiral polarity with line-shapes of CD originating from the orientational geometry variation at different bit spots (Figure 3B),. Initially, a single peak/dip response was observed when the AuNRs were oriented at small angles less than 15°, forming nearly parallel displaced geometry (*particle a*). As shown in experimental measurements (Figure 3C), the polarity from single dips correspond to down-spin as 1-bit and up-spin as 0-bit. EWFD simulations of two AuNRs arranged at ∼15° was performed (**Figure S15**), which reproduce this feature, revealing optically bright antibonding mode, and a nearly negligible optically dark anti-parallel dipole. In twisted nanorod dimers, the bonding mode has a lower energy but is optically dark because its dipoles nearly cancel out at small angles, resulting in a weak transition moment. Conversely, the antibonding mode has higher energy due to charge repulsion yet is optically bright since its dipoles align in parallel and add constructively. The single peak/dip behavior was also observed when the AuNRs formed a slightly open V-shape as shown in *particle d*. EWFD confirms this bonding and antibonding mode have closed energy level and overlap with each other (**Figure S15)**, since symmetry breaking in these structures is increased so that the coupling between two nanorods get weaker and bonding and antibonding degenerate at same energy level.

In other cases, by our DNA origami design, the bisignate (double peak) CD profiles were obtained from dimers (*particle b and c*) with large angles where both bonding and antibonding plasmon modes become optically allowed in these geometries, producing a characteristic doublet spectrum (Figure 3C). So, the antibonding mode carries the same handedness as the dimer structure, while the bonding mode has opposite handedness, causing their circular dichroism (CD) signals to have opposite signs^39^. The energy separation between these modes was governed by interparticle coupling and nanorod dimensions. Nevertheless, regardless of all three dimer configurations, it will not change the polarity of CD based on left-handed and right-handed AuNR dimers. The experimental polarity spectrum signal-noise ratio is 5 +/- 1% to 9 +/- 1%. The random noise can reduce the apparent magnitude slightly, but it is negligible probability to invert the sign. Since bit encoding relies only on overall polarity, not precise amplitude or shape details. This validates that our experimental readout is reliable despite real-world noise or fabrication variability. Thus confidently, a positive CD polarity is defined as a 1-bit, while a negative CD polarity is defined as a 0-bit.

This binary classification enables programmable information encoding across large-scale arrays, where individual metamaterial bits appear indistinguishable under dark-field imaging yet can be uniquely identified through their chiroptical signatures.

### Chiral Metamaterial Bits Array

Random information distribution plays an important role in securing data authentication and maintaining data variability. After identifying two-bit distribution in the chiral metamaterials on surface as previously described, we combined optical imaging (first layer) with chiroptical bitwise information (second layer) for constructing and mapping AuNR dimer metamaterials as random access bit arrays.

We developed a two-stage chiral bit system capable of hierarchical information encoding. In the first stage, chiral metamaterials are randomly deposited on the substrate, generating a unique spatial distribution independent of neighboring structures. The second stage introduces an authentication layer based on bit assignment, where each chiral metamaterial is optically defined as “0” or “1” through its CD polarity. This dual-layer design preserves visual indistinguishability under standard optical inspection, ensuring that no information can be inferred from appearance alone, thereby providing robust protection against duplication or forgery. Furthermore, the micron-scale spacing between bits minimizes optical crosstalk and signal noise, ensuring reliable readout without the need for complex fabrication methods or isolated steps.

**Figure 4A**, **B** shows the one-bit encoding chiral metamaterial array, in which each spot appears almost identical due to uniform nanostructure occupancy, resulting in a visually indistinguishable dark-field (DF) image. Covering this optical image is a second hidden layer of information, decoded through single-molecule CD measurement. Two representative regions (Regions 1 and 2) were selected for detailed illustration and analysis. Figure 4C outlines the information retrieval workflow. The process begins with localized optical measurements of each spot’s scattering response under left- and right-circularly polarized light (LCP and RCP). The resulting CD spectra are assigned a binary value (“0” or “1”) to each position based on the line shape polarity. Representative CD spectra from region 1 are shown in **Figure S14**, where each bit is assigned according to its CD polarity and profile. Collecting single-particle spectra across the surface enables statistical analysis of 0- and 1-bit distributions and this pattern is designed and encoded by specific arrangement of chiral metamaterials on surface, resulting in a unique code for the designer. The resulting information bits suggested high degree of randomness in the encoding process, an essential feature of strongly encrypted information bits. Shannon entropy for region 1 resulted in ∼50% probability of two bits in 1 or 0 state displaying that the system is truly random.

**Figure 4.**
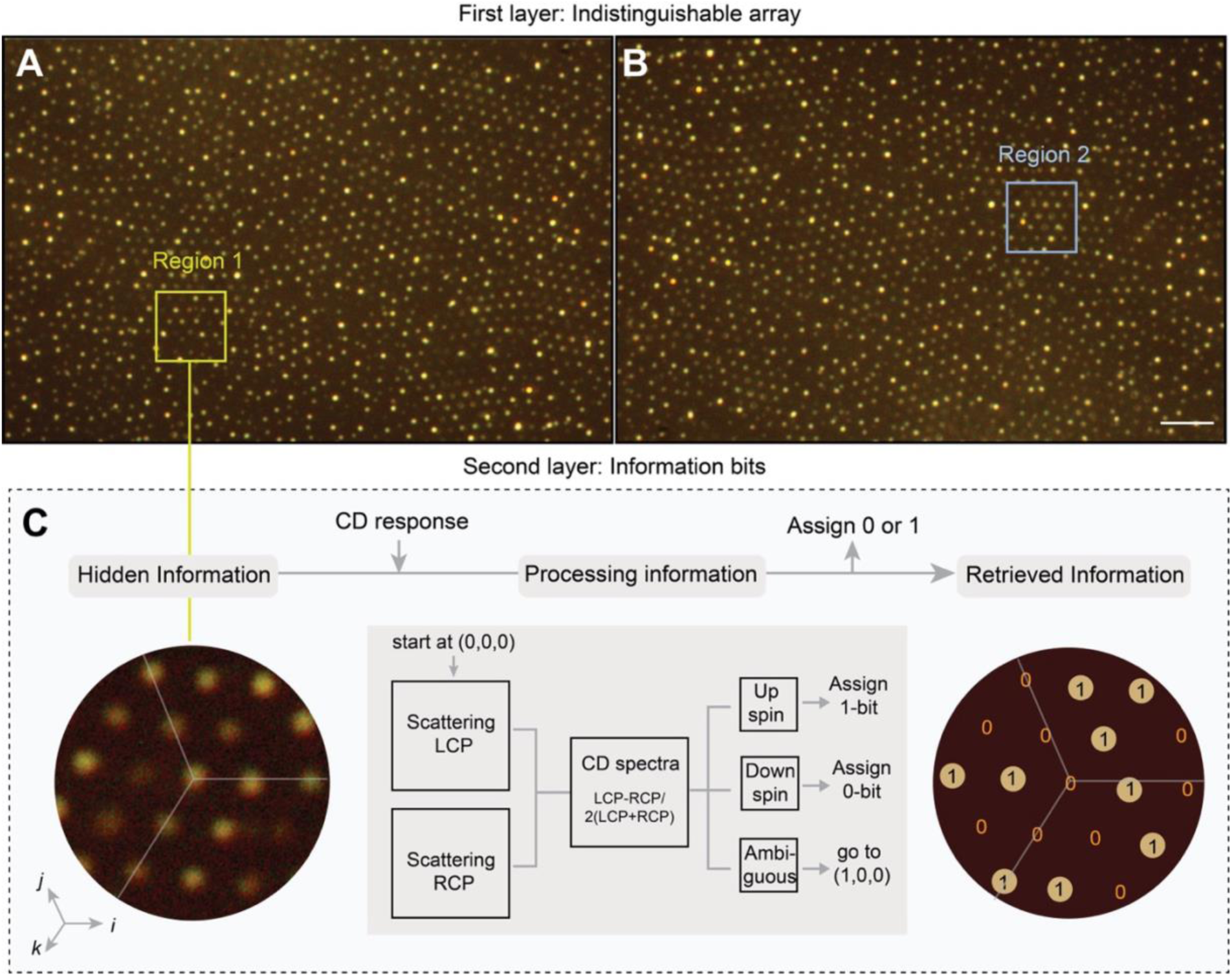
Chiral metamaterial array as random information bits. (A, B) Largescale dark-field image of chiral metamaterial array shows first layer of authentication in terms of indistinguishability of optical signals. (C) Zoomed-in region 1 from the array displaying conversion of unique CD response of each metamaterial to 0- or 1-bit based on their CD line shape adds a second layer of authentication making the system random and hard to replicate. Scale bar is 2.5 µm.

Overall, the DNA-origami templated chiral metamaterials were represented as random information bits similar to digital metamaterial arrays but capable of signal retrieval in visible light. The future is to enhance fabrication precision to enable deterministic multi-bit patterns^58^. This could involve advanced substrate engineering, where specific chiral dimers are pre-designed and positioned within a randomized array using sequence-specific DNA hybridization or site-directed origami folding, allowing for the encoding and decoding of meaningful information. Such targeted assembly would facilitate error-corrected readout protocols, potentially integrating machine learning algorithms to interpret CD maps^58^ from imaging.

Furthermore, the high modularity of the DNA origami platform opens avenues for coupling with programmable bits, such as stimuli-responsive elements (e.g., light-switchable chirality) to create reconfigurable^59,60^ arrays. To scale toward practical applications, efforts should focus on increasing storage density through denser origami lattices or multilayer stacking, aiming for gigabit-per-square-centimeter capacities while maintaining optical fidelity. Integration with complementary technologies, like quantum dot emitters^62^ or metamaterial surfaces^61^, could enhance signal-to-noise ratios and enable multi-spectral encoding for higher information depth. Ultimately, these advancements could evolve the platform into a hybrid photonic-secure system, with rigorous testing against environmental perturbations and cryptographic vulnerabilities to ensure real-world applications in fields like data security and archival storage.

## Conclusion

By integrating DNA origami-templated chiral plasmonic metamolecules with bottom-up on-surface assembly, this work establishes a robust and versatile platform for visible-range digital metamaterials. Individual gold nanorod dimers embedded in 3D chiral architectures function as physically defined information bits whose binary states are encoded in the polarity of their single-particle CD response, hidden under conventional intensity or color-based imaging. The SiMPLE patterning enables parallel large-area fabrication with long-term structural and optical stability after silicification. The resulting chiral metamaterial arrays offers two complementary security layers, a macroscopic dark-field pattern and a nanoscale addressable binary code. By information encoding into chiral metamaterial and retrievable through single-molecule CD, this approach can be translated into anti-counterfeiting labels and physical unclonable functions.

### Experimental procedures Resource availability Lead contact

Further information and requests for resources should be directed to the lead contact, Sui Yang and Hao Yan.

## Methods

### Chiral Metamaterial Fabrication

All DNA strands were purchased from Integrated DNA Technologies Inc. Probe strands (with single stranded overhangs for binding with single-strand DNA conjugated AuNR) and thiol containing strands (for conjugation with AuNRs) were purified by denaturing polyacrylamide gel electrophoresis before use. Staple strands for the DNA origami were used without further purification. The AuNRs were purchased from nanopartz and conjugated with ssDNA in-house. Single stranded scaffold strand containing 7249 nucleotides from M13m18 viral DNA was purchased from tilibit.

### DNA origami design, assembly and purification

Detailed design and sequences are shown in **Figure S1** and **Table S3**. Triangle origami was assembled by mixing 10 nM scaffold strand with the staple strands at a 1:10 molar ratio, and the designated top and bottom face probe strands (each carrying the 15-nt overhang for binding with the AuNR conjugated ssDNA strand) also at a 1:10 molar ratio, in 1xTAE-Mg buffer (12.5 mM). Thermal annealing was followed, from 65 °C to 4 °C in ∼ 18 h:^38^ 65 °C at a rate of 1 °C per 15 min, 64 °C to 60 °C at a rate of 1 °C per 5 min, 59 °C to 40 °C at a rate of 1 °C per 45 min, 39 °C to 36 °C at a rate of 1 °C per 30 min, 35 °C to 20 °C at a rate of 1 °C per 5 min, then hold at 4 °C.

The solution was put into a 100 kDa molecular Amicon centrifugal filter and washed with 1xTAE-Mg, 5 times to remove access staple strands. The purified samples were stored at -20 °C until further use.

### Conjugation of single strand DNAs with AuNR

AuNRs (60x20 nm) were conjugated with top and bottom ssDNA handles using low pH method.^42^ Orthogonal ssDNA handles were synthesized in-house on solid-phase DNA synthesizer and HPLC purified before use. ssDNA functionalized AuNRs were purified by centrifugation with 0.1% SDS in 0.5x TBE at 5000 rpm for 20 mins, 3 times to remove excess probe strands. Pelleted AuNRs were resuspended in the supernatant to a concentration of ∼ 5 nM for subsequent conjugation with origami.

### Functionalization, purification and characterization of chiral metamaterials

The purified triangle origami containing overhangs was mixed with top and bottom ssDNA-AuNR conjugates in a 1:3 molar ratio in 10 mM MgCl_2_. Thermal annealing was followed for ∼13 h from 45 °C to 25 °C at a rate of 1 °C per min for 4 repeated cycles. A 0.75% agarose gel electrophoresis (1.5 hr at 100 V) in 1xTAE-Mg was used to purify the final triangle containing both top and bottom AuNR constructs, desired conjugate band was excised, followed by smashing in a spinX tube. This sample was kept in –20° C for 1 hr before centrifuging it down at 5000 rpm for 10 mins. Pelleted triangle-AuNRs construct was resuspended, and concentration of the sample was calculated using nucleic-acid A(260) value. These conjugates were stored at –20° C until use.

Note: Leaving them too long over one month can lead to aggregation and can lead to a difference in single occupancy on nanoarray as discussed later. About 5 chips of chiral array can be made without aggregation when starting with 200 uL, 5 nM AuNR.

The purified chiral metamaterials were characterized for yield using Transmission electron microscopy (TEM). 5µl of each purified sample (2 nM) was applied to a commercially supplied Formvar-stabilized carbon type-B, 400mesh copper grids (Ted Pella, part number 01814-F) that had been glow discharged for 1 min at 15mA using a Pelco easiGlow glow-discharge system (Ted Pella, Redding, CA, USA), and stained using 5µl of a freshly prepared 2% aqueous uranyl formate solution containing 25 mM sodium hydroxide (NaOH). Samples were incubated for 60 to 90 s depending on sample conditions. Excess liquid was wicked away with Whatman filter paper 1, and grids were left to dry for 30-60 min prior to imaging. Images were acquired on a Talos microscope (Thermo Fisher LMT) operated at 120 kV accelerating voltage, using a charge-coupled device (CCD) camera at 73000x magnification.

### Silica Microsphere assisted Pattering via Liquid Elimination (SiMPLE)

Fabrication of binding spots using Silica microsphere assisted pattering via liquid elimination (SiMPLE), DNA triangle origami placement and chiral metamaterial array formation.

Considering, ∼1 um size distribution for silica and polystyrene, we observed the contact areas generated by silica microspheres is 7.9x10^3^ nm² and polystyrene nanoparticles is 2.5x10^4^ nm². All measurements are done using Bruker ScanAsyst AFM in air-mode.

### Materials and equipment required

1. 10×10 mm^2^ coverslips (Ted Pella, 260375-15).
2. Plasma cleaner (Harrick Basic Plasma Cleaner PDC-32G/PDC-3°2G-2)
3. Oven, Hotplate and stirrer (Denville)
4. Desiccator (Hach, Product no. 223830)
5. Branson ultrasonic bath, AFM (Bruker FastScan)
6. Appropriately sized Silica microspheres (NanoXact, 1um, Nanocomposix)
7. Passivation agent: HMDS (440191–100 mL, Sigma)
8. Plastic Petri dish 35 mm (Corning)
9. N,N-Dimethylformamide anhydrous (DMF), 99.8% (CAS: 68-12-2, Sigma)
10. Dichloromethane (DCM) (Sigma)
11. MilliQ water

### Protocol for binding site fabrication

1. Wash glass slides with isopropanol (IPA) for 2 min and blow-dry with nitrogen
2. Mark the slides on the bottom with a glass cutter to locate area for colloidal mask deposition
3. Perform a 10–min air plasma cleaning in Harrick plasma cleaner at ∼18 W (“High” setting) to activate the glass surface
4. In an Eppendorf tube, weigh 35 mg of 1 µm silica nanoparticles and add 60 µL of DMF. Sonicate for 10 mins. In a new 200 µL tube, pipette 14 µL of Silica-in-DMF stock and add 6 µL of DCM and sonicate for 10 mins
5. Place the activated glass slide in a plastic petri dish (dia, 35 mm) and submerge in ∼1 mL of MilliQ water.
6. Gently drop ∼1-2 µL of silica suspension made in step 4 on water. A silica film will form on the air/water interface as the particles pack closely.
7. Carefully remove the water from the dish with a pipette, taking care not to introduce vibrations. This creates a self-assembled monolayer slowly on the surface of the glass slide.
8. Transfer the coated coverslip to a desiccator and place in oven at 150 °C for 15 mins to remove left-over moisture. Once dried, a diffraction pattern can be observed, confirming the existence of a close-packed monolayer/multilayer of nanospheres.
9. Carry out a 5 min “descum” plasma in air at ∼18 W in Harrick plasma cleaner on the monolayer.
10. Transfer the coverslip to a desiccator, add 8–10 drops of HMDS (in a glass cuvette), and deposit under a vacuum seal for 1 hr.
11. Lift-off silica nanospheres with water sonication in a Branson ultrasonic bath for 30–60 sec and blow dry with a nitrogen gun to create origami binding sites.
12. Bake at 120 °C for 5 min to stabilize the HMDS on the surface.

### Protocol for DNA origami placement

1. Rinse the chip with NiCl_2_ (1 mM) in Tris-HCl (40 mM) buffer (pH ∼ 6.5) for 2 mins.
2. Incubate chips with appropriate concentration of origami triangle ∼100-200 pM diluted in 40 mM Tris-HCl and 40 mM MgCl_2_, pH 8.3 for 1 hr.
3. Wash in containing ∼40 mM MgCl_2_, 40 mM Tris-HCl, pH ∼ 8.3 in a petri dish for 5 min using a low-speed orbital shaker. Repeated aspiration of buffer also works equally well.
4. Transfer the chip to petri dish containing ∼40 mM MgCl_2_, 40 mM Tris-HCl + 0.07% tween 20, pH ∼ 8.4 for another 5 mins.
5. Lastly, wash with ∼35 mM MgCl_2_, 10 mM Tris-HCl, pH ∼8.9 to hydrolyze HMDS and remove any non-specifically bound origami in the background.
6. For AFM characterization, transfer to ethanol drying series: 10s in 50% ethanol, 20s in 75% and 2min in 85% ethanol.

### Protocol for Chiral metamaterial placement

1. Follow all steps for binding site fabrication and washing steps as for origami only above.
2. Chiral metamaterials are directly incubated on array for 1 hr at required concentrations (120, 150, 200 pM diluted in 40 mM Tris-HCl and 40 mM MgCl_2_).

Note: Chiral metamaterial (60x20 nm) required a higher concentration (250 pM) for complete deposition than smaller chiral metamaterials due to possibility of settling down and aggregation of the large AuNRs. These aggregations seemed to be less when working with smaller (38x10 nm) AuNR, where same concentration as only origami is required. For high quality chiral metamaterial array, the fresh chiral metamaterials should be prepared before deposition on array.

### DNA-PAINT and Photobleaching Experiments

All total internal reflection fluorescence (TIRF) experiments utilized a benchtop super-resolution Oxford Nanoimager (Oxford, UK). After three washes with buffer, chip containing origami with overhangs for imager was stacked on double-sided Kapton tape to form a flow chamber. Imager solution, containing up to 5 nM oligonucleotides along with an oxygen scavenging mix (comprising 2x PCA, 3x PCD, and 5x Trolox-Quinone), was introduced into the chamber and left to incubate for one hour. DNA-PAINT data set was collected for 20 minutes and analyzed using Picasso. The movie files were cropped to 256 by 256 pixels using FIJI and then processed movie was imported into the Picasso Localize module to generate precise localization maps using defined parameters. To achieve high-quality super-resolution images, the Picasso Render module applied redundant cross-correlation (RCC) for drift correction, while outlier localizations were filtered by x/y position precision and background noise were excluded using the Picasso Filter module. All processing steps were carried out on an Alienware desktop fitted with an Intel Core i7-6800K CPU, 32 GB RAM, and an NVIDIA GeForce GTX 1080 GPU.

For photobleaching experiments, a similar flow chamber as described above was prepared. DNA origami with overhangs for the photobleaching imager strand was with photobleaching-imager strand in 20x excess with cy3b along with oxygen scavengers. Laser power was carefully modified to produce a gradual bleaching gradient, facilitating the identification and counting of fluorescence steps. Quantification was performed using two software approaches: ImageJ and iSMS as described previously. For the latter approach, the field of view was cropped, and both regions of interest (ROIs) were aligned to enable accurate step counting.

### Single-molecule Circular Dichroism (CD) measurements Optical set-up for single molecule CD measurements

Colocalization technique was applied to locate the chiral structured AuNR on glass substrates under SEM (Zeiss-Auriga). The same chiral structures were imaged with free-space dark field scattering setup. The dark field scatterings were recorded from different structures. To capture scattering signal from single metamaterial, a free space dark-field microscope was assembled with point-to-point measurement setup shown below. Unpolarized light from HAL (halogen) light source passed through linear polarizer and quarter (¼) waveplate subsequently to become circularly polarized light. The relative angle between linear polarizer and ¼ waveplate determine the chirality of polarized light for LCP at -45° and RCP at 45°. Left chiral polarized light as LCP: Ex=E0*exp(-j*ewfd.k0*z), Ey=-i*E0*exp(-j*ewfd.k0*z and right chiral polarized light as RCP: Ex=E0*exp(-j*ewfd.k0*z), Ey=-i*E0*exp(-j*ewfd.k0*z). With scattering boundary condition, LCP and RCP excited single chiral structure. Due to selection rule of photonic energy states, bonding states and anti-bonding states corresponding to specific structures, which lead to different absorptions.

To collect the absorption difference, top dark filed oil-immersion lens introduces light at high angle with index of 1.4 immersion oil. 100X objective collected scattering signal that was collected by the OMAX A35180U3 for optical images of array. Andor spectrometer (SR-303i-B with grating 150/500) was used to collect scattering signal.

Circular Dichroism Scattering was calculated using this equation:

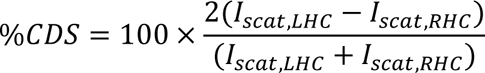

### Electromagnetic Waves, Frequency Domain (EWFD) Calculations

Scattering-field electromagnetic simulations were performed using COMSOL Multiphysics with the electromagnetic waves, frequency domain (EWFD) module in 3D. The plasmonic dimer system was modeled as two gold nanorods with a radius of 30 nm and length of 60 nm, separated by a fixed inter-rod distance of 2 nm. The surrounding effective medium was defined with a constant refractive index of n = 1.33 to approximate the effective dielectric environment. The simulation domain was enclosed within perfectly matched layers (PMLs) with a thickness of 38 nm to eliminate spurious reflections from boundaries. Circularly polarized lights (left- and right-circularly polarized, LCP and RCP) were incident normal to the dimers to excite the system. Scattering spectra were computed over a wavelength range from 400 to 1000 nm. The differential scattering spectra (CD_scat_) were obtained by subtracting the scattering intensities under LCP and RCP illumination as CD_scat_ = LCP_scat_ -RCP_scat_).

## Acknowledgements

We would like to thank Axel Whittman for helping with carbon coating. We acknowledge the use of SEM and TEM within the Eyring Materials Center at Arizona State University. The research reported in this project has received funding support from the National Science Foundation SemiSynBio III grant no. 2227650 to H.Y., S. Y., R. H. and P. S.

## Authors Contribution

S. Y., H.Y., R. H., P. S., conceptualized the idea. D. S. and S. F. wrote the manuscript, D.S., S. F., P. C., designed, performed and interpreted all experiments. D. S., P. C., L. W., N. B. and A. S., prepared SiMPLE array and performed all troubleshooting on fabrication, D. S., S. F., P. C., A.P. and L. W., performed high-resolution and optical imaging, D. S., S. F., S. R. and L. C. performed EWFD and oxDNA simulations. S. Y., H.Y., R. H. and P. S. oversaw the whole project, mentored and provided guidance. All authors edited the final version of manuscript.

## References

1. Ouyang, X. et al. Synthetic helical dichroism for six-dimensional optical orbital angular momentum multiplexing. Nat. Photon. 15, 901–907 (2021).

2. Han, M., Gao, X., Su, J. Z. & Nie, S. Quantum-dot-tagged microbeads for multiplexed optical coding of biomolecules. Nature 19, 631–635 (2001).

3. Organick, L. et al. Random access in large-scale DNA data storage. Nat. Biotechnol. 36, 242–248, (2018).

4. Goldman, N. et al. Towards practical, high-capacity, low-maintenance information storage in synthesized DNA. Nature 494, 77–80, (2013).

5. Zhang, Y. et al. Information stored in nanoscale: Encoding data in a single DNA strand with Base 64. Nano Today 33, 6–11 (2020).

6. Wiecha, P. R. et al. Evolutionary multi-objective optimization of color pixels based on dielectric nanoantennas. Nat. Nanotechnol. 12, 163–169 (2017).

7. Matoba, B. O., Nomura, T., Pe, E. & Javidi, B. Optical Techniques for Information Security. Proc. IEEE. 97, (2009).

8. Psaltis, D. Coherent Optical Information Systems. Science 298, 1359–1363 (2002).

9. Bellare, M., Kilian, J., Rogaway, P. The security of the cipher block chaining message authentication code. J. Comput. Syst. Sci. 61, 362–399 (2000).

10. Capkun, S., et al. Integrity Codes: Message Integrity Protection and Authentication over Insecure Channels. IEEE TDSC 5, 208–223 (2008).

11. Kim, J. H. et al. Nanoscale physical unclonable function labels based on block co-polymer self-assembly. *Nat*. Electron. 5, 433–442 (2022).

12. Wu, J. et al. A High-Security mutual authentication system based on structural color-based physical unclonable functions labels. Chem. Eng. J. 439, 135601 (2022).

13. Gao, Y., Al-Sarawi, S. F. & Abbott, D. Physical unclonable functions. Nat. Electron. 3, 81–91 (2020).

14. Giovampaola, C. Della, & Engheta, N., Digital metamaterials. Nat. Mat. 13, 1115–1121 (2014).

15. Cui, T. J., Liu, S., Bai, G. D. & Ma, Q. Direct Transmission of Digital Message via Programmable Coding Metasurface. Research. 2019, (2019).

16. Dai, J. Y. Independent control of harmonic amplitudes and phases via a time-domain digital coding metasurface. Light Sci. Appl. 7, (2018).

17. You, A., Be, M. & In, I. A spatial light modulator for terahertz beams. Appl. Phys. Lett. 94, 213511 (2009).

18. You, A., Be, M. & In, I. An electrically driven terahertz metamaterial diffractive modulator with more than 20 dB of dynamic range. Appl. Phys. Lett. 104, 091115 (2014).

19. Gao, L. et al. Broadband diffusion of terahertz waves by multi-bit coding metasurfaces. Light Sci. Appl. 4, 324 (2015).

20. Wang, Z., Zhang, Q., Zhang, K. & Hu, G. Tunable Digital Metamaterial for Broadband Vibration Isolation at Low Frequency. Adv. Mat. 28, 9857–9861 (2016).

21. Xie, B. et al. Coding Acoustic Metasurfaces. Adv. Mat. 29, 1–8 (2017).

22. Ahamed, E., Faruque, M. R. I., Alam, M. J., Mansor, M. F. Bin & Islam, M. T. Digital metamaterial filter for encoding information. Sci. Rep. 10, 2–10 (2020).

23. Zhang, L., Liu, S., Li, L. & Cui, T. J. Spin-Controlled Multiple Pencil Beams and Vortex Beams with Di ff erent Polarizations Generated by Pancharatnam-Berry Coding Metasurfaces. ACS Appl. Mater. Interfaces. 9, 36447–36455 (2017).

24. Cui, T. J., Qi, M. Q., Wan, X., Zhao, J. & Cheng, Q. Coding metamaterials, digital metamaterials and programmable metamaterials. Light Sci. Appl. 3, 218–218 (2014).

25. Engheta, N. Tunneling of Electromagnetic Energy through Subwavelength Channels and Bends using Near-Zero Materials. Phys. Rev. Lett. 97, 1–4 (2006).

26. Shen, Z. et al. Design of transmission-type coding metasurface and its application of beam forming. Appl. Phys. Lett. 109, 121103, (2016).

27. Liu, S. & Cui, T. J. Concepts, Working Principles, and Applications of Coding and Programmable Metamaterials. Adv. Opt. Mater. 5, 1–27 (2017).

28. Paul, A. & Timp, G. How to Manufacture Photonic Metamaterials. *Adv*. Mater. Technol. 10, 1–27 (2025).

29. Bahk, Y. M., Kim, D. S. & Park, H. R. Large-Area Metal Gaps and Their Optical Applications. Adv. Opt. Mater. 7, 1–21 (2019).

30. Liu, M., Powell, D. A., Shadrivov, I. V., Lapine, M. & Kivshar, Y. S. Spontaneous chiral symmetry breaking in metamaterials. Nat. Commun. 5, 1–9 (2014).

31. Avetisov, V. & Goldanskii, V. Mirror symmetry breaking at the molecular level. Proc. Natl. Acad. Sci. U. S. A. 93, 11435–11442 (1996).

32. Qiu, M. et al. 3D Metaphotonic Nanostructures with Intrinsic Chirality. Adv. Funct. Mater. 28, 1–16 (2018).

33. Kotov, N. A., Liz-Marzán, L. M. & Weiss, P. S. Chiral Nanostructures: New Twists. ACS Nano 15, 12457–12460 (2021).

34. Fernandez-Corbaton, I. et al. New Twists of 3D Chiral Metamaterials. Adv. Mater. 31, 1–7 (2019).

35. Halas, N. J., Lal, S., Chang, W. S., Link, S. & Nordlander, P. Plasmons in strongly coupled metallic nanostructures. Chem. Rev. 111, 3913–3961 (2011).

36. Schellman, J. A. Flexibility of DNA. Biopolymers 13, 217–226 (1974).

37. Prodan, E., Radloff, C., Halas, N. J. & Nordlander, P. A Hybridization Model for the Plasmon Response of Complex Nanostructures. Science. 302, 419–422 (2003).

38. Eyring, H., Liu, H. C. & Caldwell, D. Optical rotatory dispersion and circular dichroism. Chem. Rev. 68, 525–540 (1968).

39. Model, B. K., Yin, X., Scha, M., Metzger, B. & Giessen, H. Interpreting Chiral Nanophotonic Spectra: The Plasmonic Born−Kuhn Model. Nano Lett. 13, 6238–43 (2013).

40. Hu, Z. et al. Plasmonic Circular Dichroism of Gold Nanoparticle Based Nanostructures. Adv. Opt. Mater. 7, 1–13 (2019).

41. Zhang, Q. et al. Unraveling the origin of chirality from plasmonic nanoparticle-protein complexes. Science 365, 1475–1478 (2019).

42. Chakraborty, A. et al. The Influence of the Supporting Substrate on Single-Particle Circular Differential Scattering of DNA Assembled Nanorod Dimers. Adv. Optical Mater. 13, 1–10 (2025).

43. Rothemund, P. W. K. Folding DNA to create nanoscale shapes and patterns. Nature 440, 297–302 (2006).

44. Kuzyk, A. et al. DNA-based self-assembly of chiral plasmonic nanostructures with tailored optical response. Nature 483, 311–314 (2012).

45. Shen, X. et al. Rolling up gold nanoparticle-dressed dna origami into three-dimensional plasmonic chiral nanostructures. J. Am. Chem. Soc. 134, 146–149 (2012).

46. Dai, G. et al. DNA origami-directed, discrete three-dimensional plasmonic tetrahedron nanoarchitectures with tailored optical chirality. ACS Appl. Mater. Interfaces 6, 5388–5392 (2014).

47. Dietz, H., Douglas, S. M. & Shih, W. M. Folding DNA into twisted and curved nanoscale shapes. Science 325, 725–730 (2009).

48. Douglas, S. M. et al. Self-assembly of DNA into nanoscale three-dimensional shapes. Nature 459, 414–418 (2009).

49. Biswas, A. et al. Advances in top-down and bottom-up surface nanofabrication: Techniques, applications & future prospects. Adv. Colloid Interface Sci. 170, 2–27 (2012).

50. Wen, B., Yang, J., Hu, C., Cai, J. & Zhou, J. Top-Down Fabrication of Ordered Nanophotonic Structures for Biomedical Applications. Adv. Mater. Interfaces 11, 1–20 (2024).

51. Chopade, P., Hariadi, R., Satyabola, D., Sakai, A. “Methods of Fabricating High-Throughput Arrays with Customizable Properties Using Silica Microsphere Assisted Patterning,”. (U.S. provisional patent application 63/722,979 (2024).

52. Gandla, S. et al. Random laser ablated tags for anticounterfeiting purposes and towards physically unclonable functions. Nat. Commun. 15, (2024).

53. Chong, C. N., Jiang, D., Zhang, J. & Guo, L. Anti-counterfeiting with a random pattern. Proc. - 2nd Int. Conf. Emerg. Secur. Inf., Syst. Technol. Secur. 146–153 (2008).

54. Caruthers, J. M. et al. Physical One-Way Functions. 297, 2026–2030 (2026).

55. Arppe, R. & Sørensen, T. J. Physical unclonable functions generated through chemical methods for anti-counterfeiting. Nat. Rev. Chem. 1, 31 (2017).

56. Dass, M. et al. Self-Assembled Physical Unclonable Function Labels Based on Plasmonic Coupling. Adv. Opt. Mater. 13, (2025).

57. Shetty, R. M., Brady, S. R., Rothemund, P. W. K., Hariadi, R. F. & Gopinath, A. Bench-Top Fabrication of Single-Molecule Nanoarrays by DNA Origami Placement. ACS Nano 15, 11441–11450 (2021).

58. Yue, W. et al. Physical unclonable in-memory computing for simultaneous protecting private data and deep learning models. Nat. Commun. 16, 1–13 (2025).

59. Kuzyk, A. et al. Reconfigurable 3D plasmonic metamolecules. Nat. Mater. 13, 862–866 (2014).

60. Schreiber, R. et al. Chiral plasmonic DNA nanostructures with switchable circular dichroism. Nat. Commun. 4, 1–6 (2013).

61. Martynenko, I. V. et al. Site-directed placement of three-dimensional DNA origami. Nat. Nanotechnol. 18, 1456–1462 (2023).

62. Luo, X. et al. DNA origami directed integration of colloidal nanophotonic materials with silicon photonics. bioRxiv 2025.01.23.634416 (2025).

