## Supplementary File for "DNA-templated Chiral Metamaterial Array as Information Bits"

### Section 1. Supplementary Figures

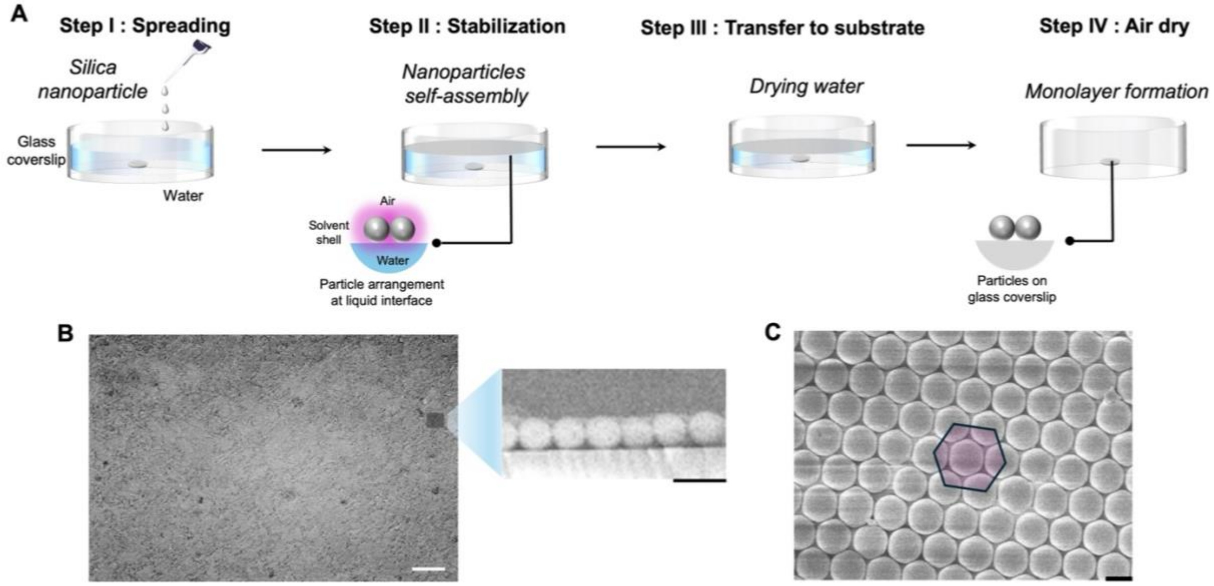


**Figure S1. Procedure of monolayer formation using SiMPLE.** (A) In step I, silica nanoparticles were added to cover the air-water interface slowly, followed by Step II, where the nanoparticles are self-assembled to stabilize them on surface, in Step III, water is slowly removed from sides , finally in Step IV, remaining water is allowed to dry on its own. (B) Left: Large area self-assembly of silica nanoparticles, right: SEM cross section of silica nanoparticles on glass substrate. (C) Top view SEM of closely packed silica nanoparticles in hexagonal packing. Scale bar of (B) is 50 µm, right: 2 µm and (C) is 1 µm

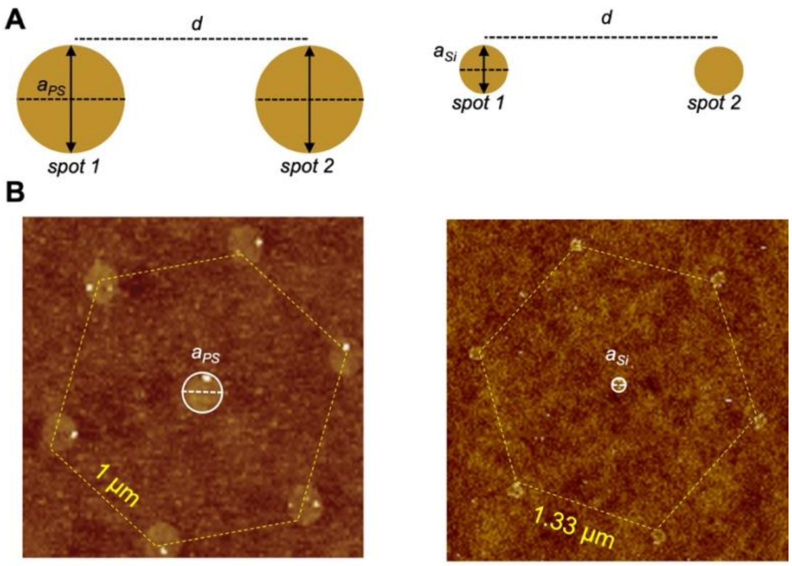


**Figure S2. Comparison of binding spots generated by polystyrene and silica beads.** Illustration of binding spots generated by polystyrene and silica beads due to differences in their stiffness and corresponding AFM images in air-mode confirms a smaller binding spot generated with silica beads.

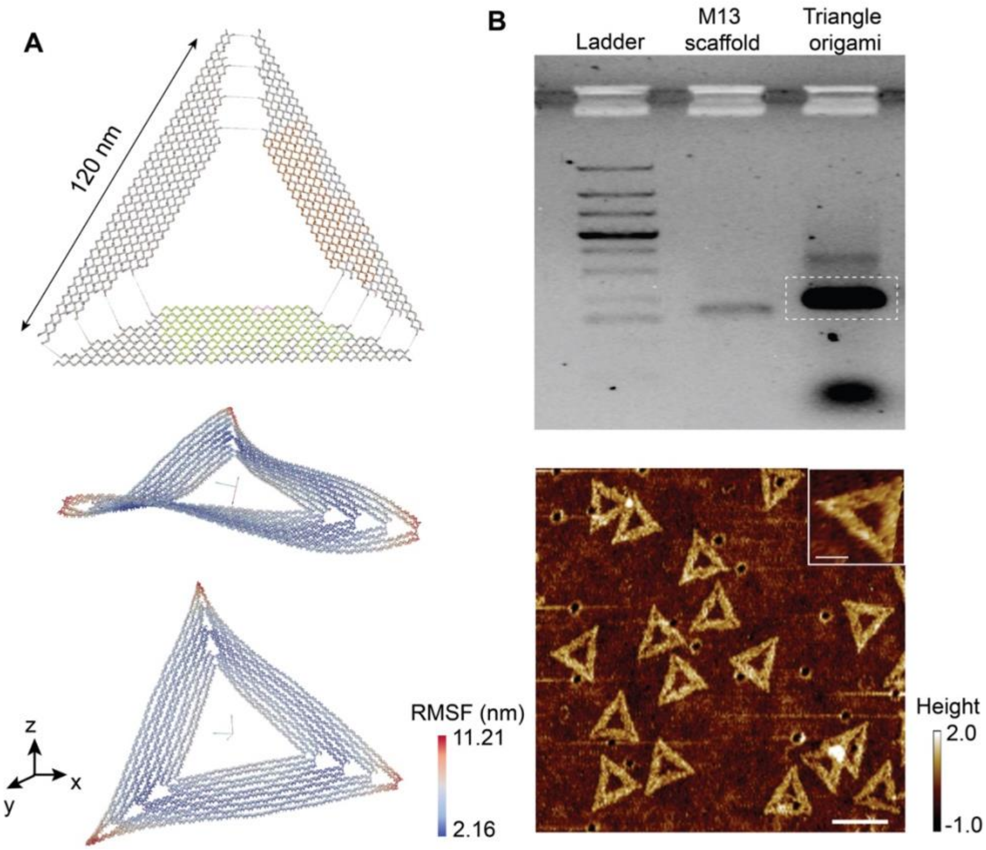


**Figure S3. Self-assembly of origami triangle.**
(A) oxDNA mean structure of symmetric DNA origami triangle. (B) 0.75% agarose gel electrophoretic mobility assay showed successful self-assembly of triangle origami nanostructures. (C) Atomic Force Micrographs (AFM) of gel purified triangle origami. Scale bar is 100 nm.

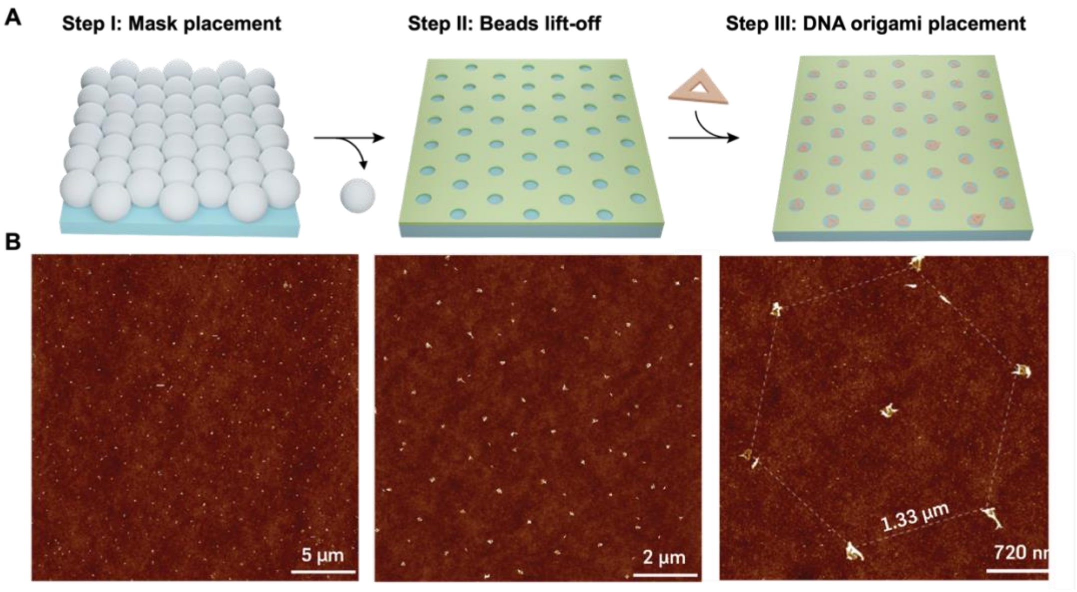


**Figure S4. DNA origami placement on SiMPLE fabricated array.** (A) Three step placement of triangle origami on array. (B) AFM images show the uniform triangle occupancy on array (slight misfolding of triangle is also visible could be due to ethanol drying).

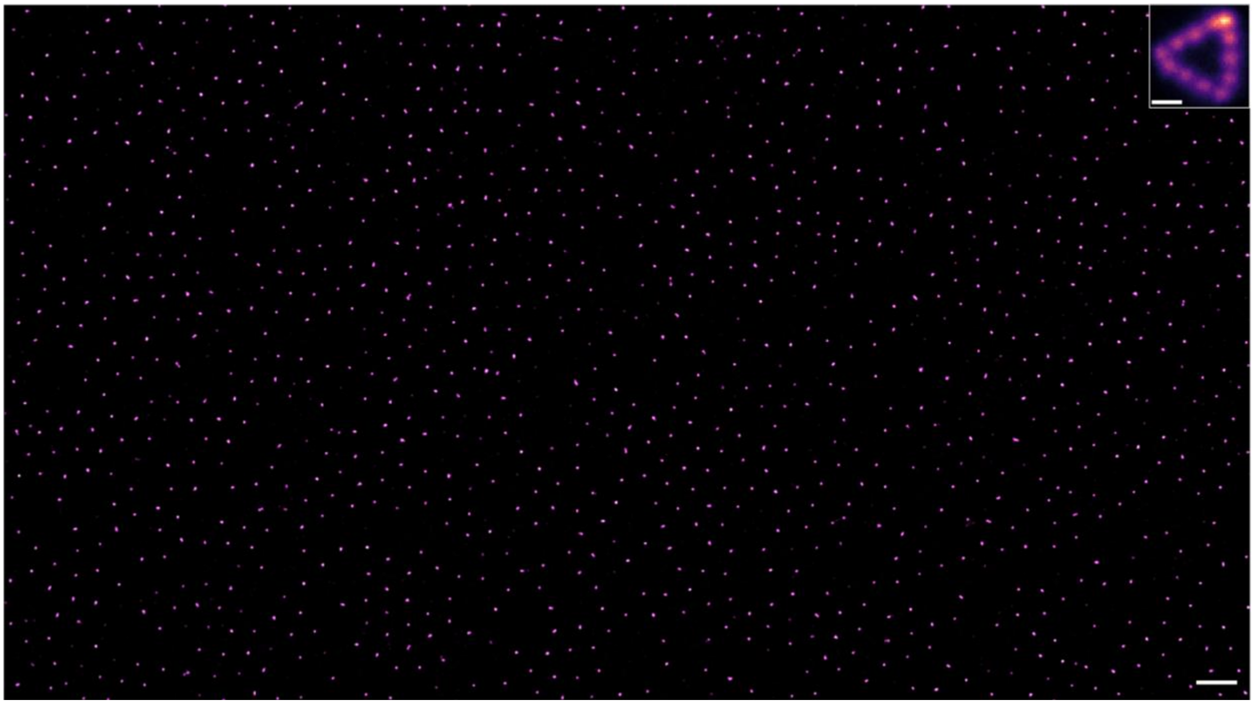

**Figure S5. DNA-PAINT showing single-occupancy of triangle origami on surface.** Scale bar of large area is 2 μm and inset is 50 nm.

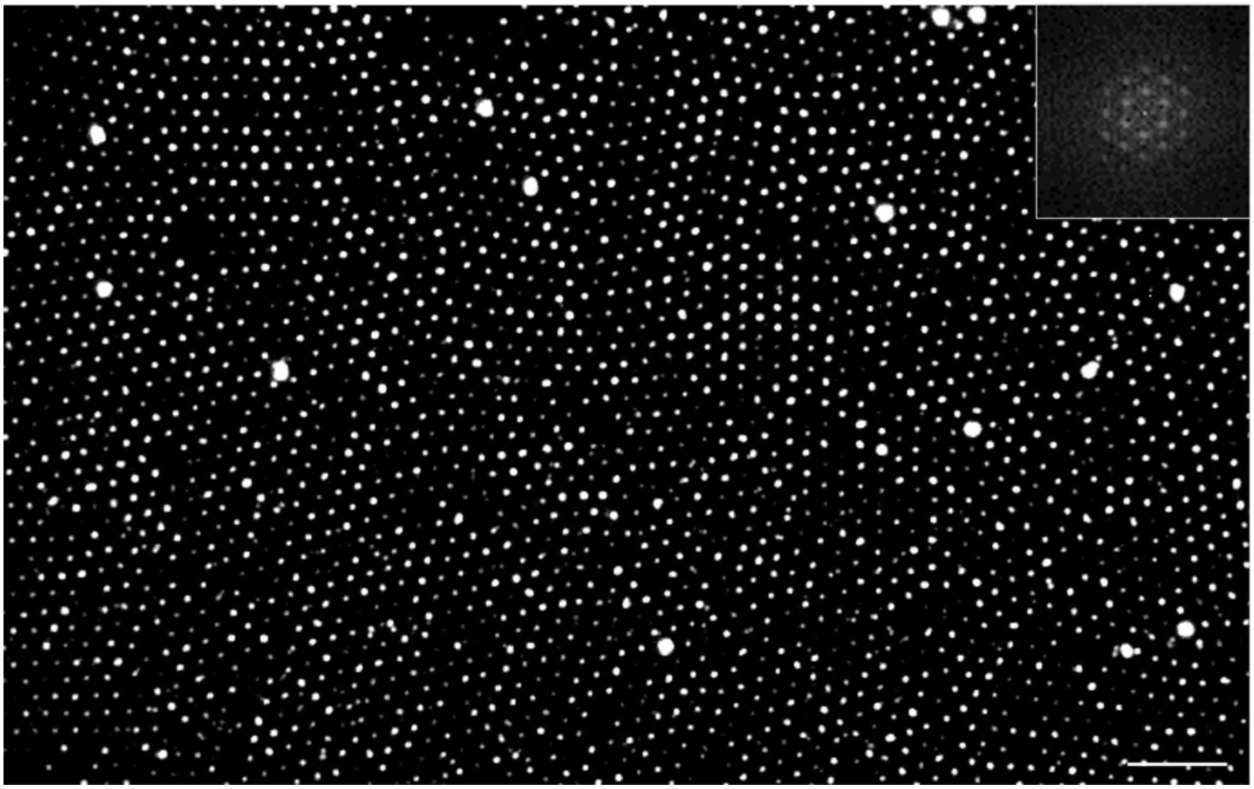

**Figure S6. Large scale placement of DNA origami triangle with cy3b imager.** Photobleaching shows a large-scale occupancy of origami on substrate, inset shows FFT image indicating a hexagonal pattern. Scale bar is 5 μm.

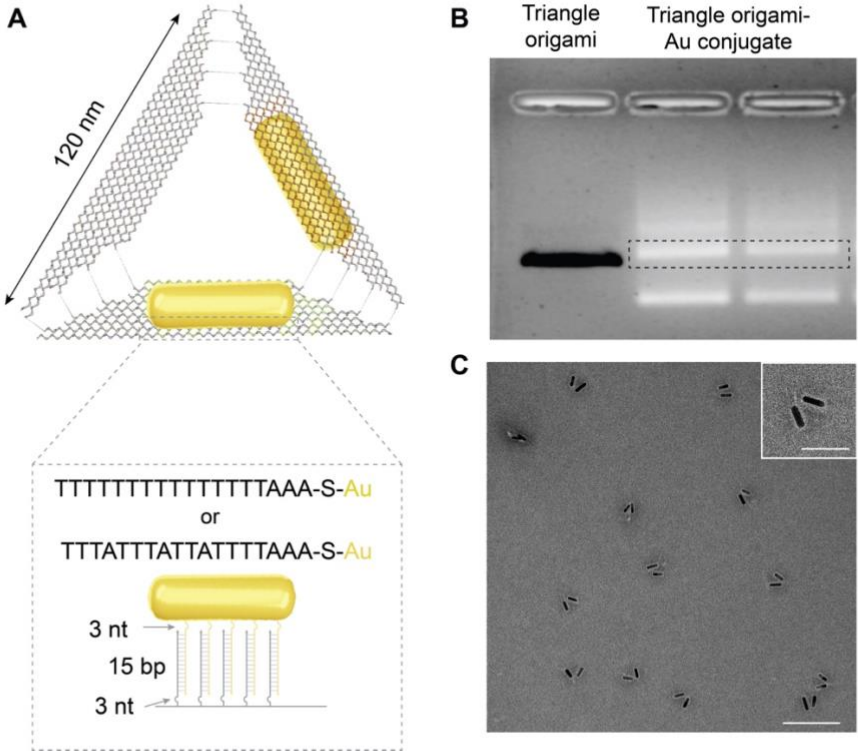


**Figure S7. Assembly of chiral metamaterials in solution.**
(A) Triangular DNA origami with specific positions for anchoring AuNR to form a symmetry breaking chiral structure. (B) 0.75% agarose gel electrophoretic mobility assay showed successful assembly of chiral metamaterials. (C) Transmission electron micrographs (TEM) of gel purified chiral metamaterials.

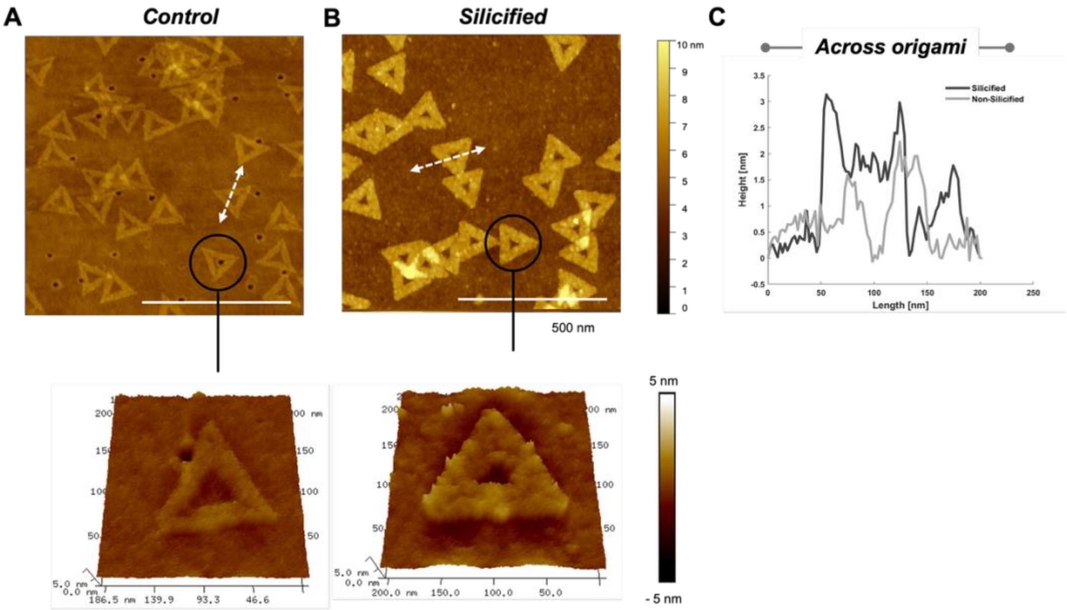


**Figure S8. Silicification of triangle origami.** (A) Triangle DNA origami control. (B) Silicified triangle origami. (C) Increase in the height of DNA origami upon silicification is evident with an increased height.

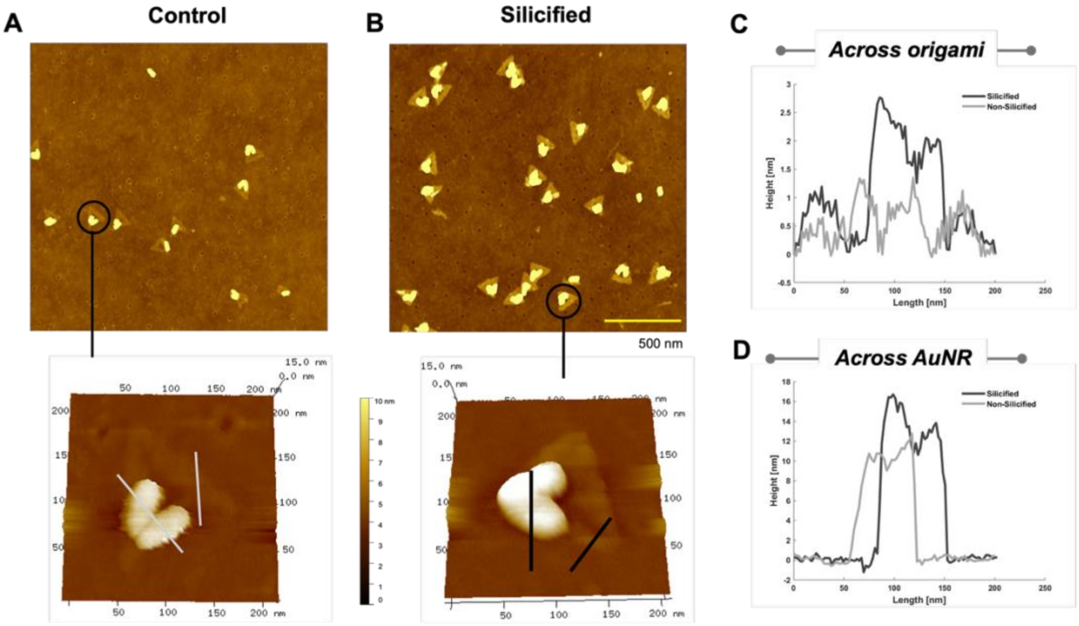


**Figure S9. Silicification of chiral metamaterials.** (A) Chiral metamaterial control. (B) Silicified metamaterials. (C) Increase in the height of DNA origami across origami and (D) across AuNR upon silicification is evident with an increased height.

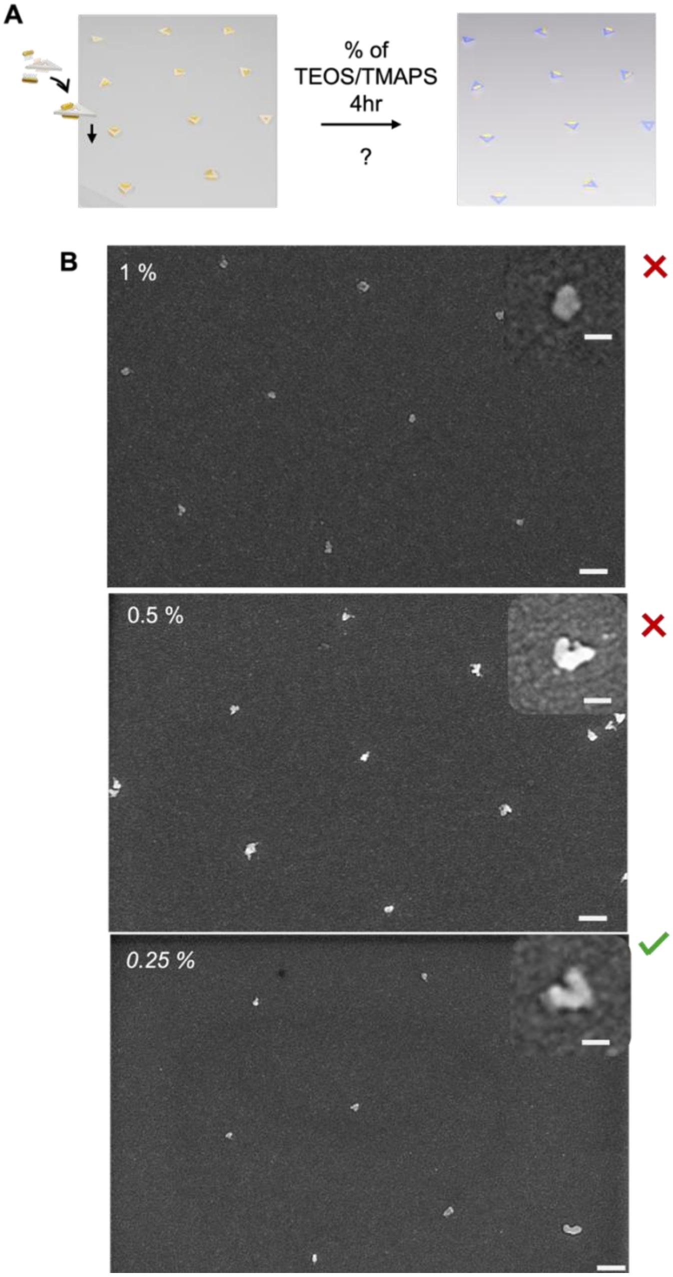


**Figure S10. Different ratios of TEOS/TMAPS used for silicification of chiral metamaterials.** (A) Silicification of chiral metamaterial deposition at different ratios of TEOS/TMAPS at 4hr. (B) Scanning electron micrographs of chiral metamaterial array at 1%, 0.5% and 0.25% of silicification solution.

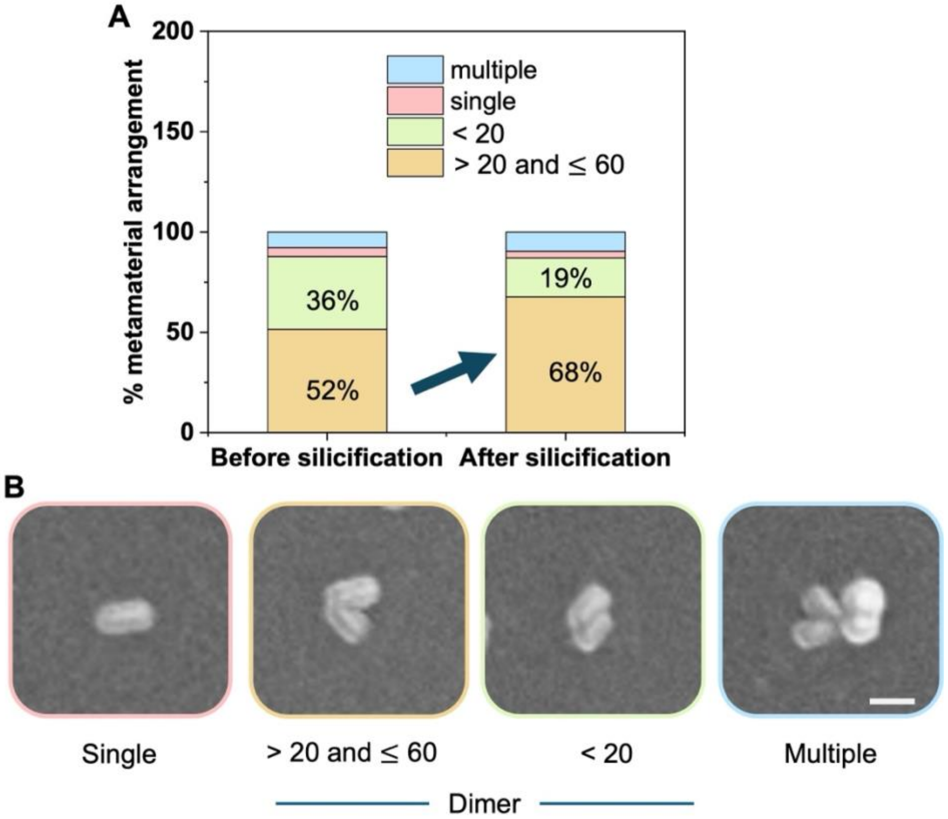


**Figure S11. Effect of silicification on various geometries of chiral metamaterials.** (A) Percentage of metamaterials before and after silicification. (B) Improvements observed in chiral arrangement upon silicification.

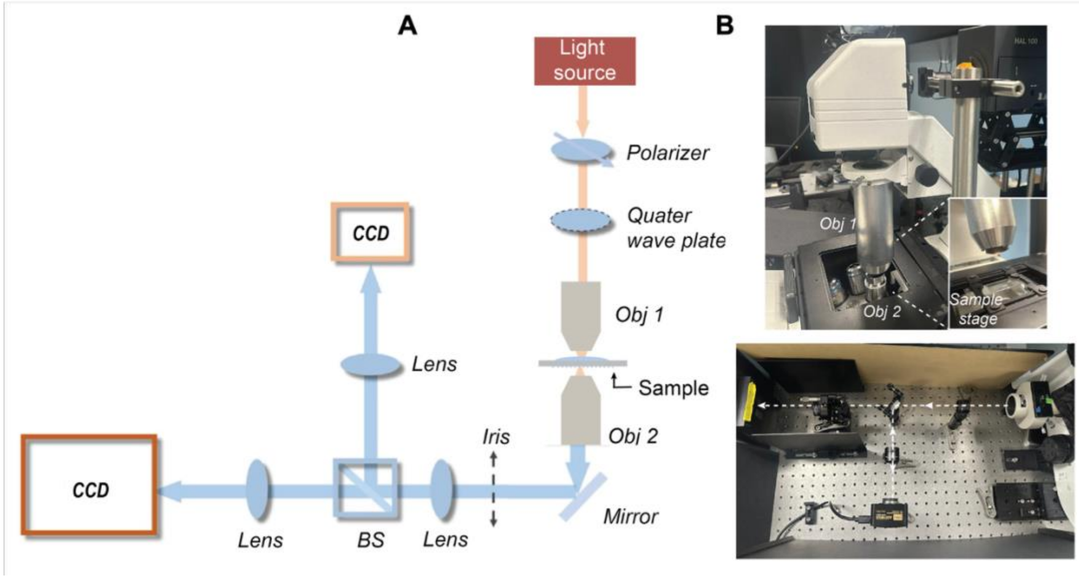


**Figure S12. Optical set-up used.** (A) Line diagram of the microscope alignment. (B) Photograph of actual set-up used in the study, white lines indicate the path of light.

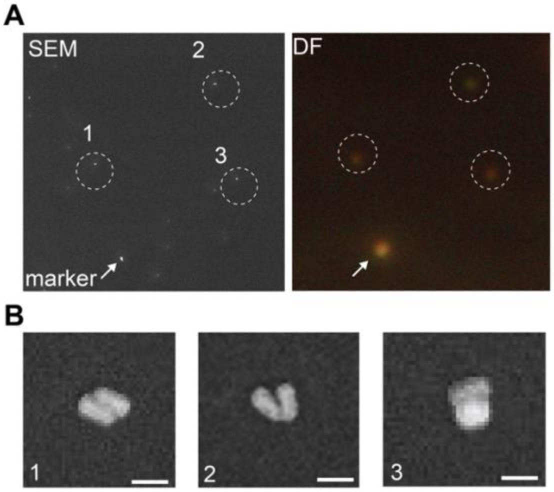


**Figure S13. Colocalization of random single metamaterials (60x20 nm).** Using a glass marker, single-molecule metamaterials were visualized under SEM and same region was identified on dark field microscope at highest exposure and gain settings on camera.

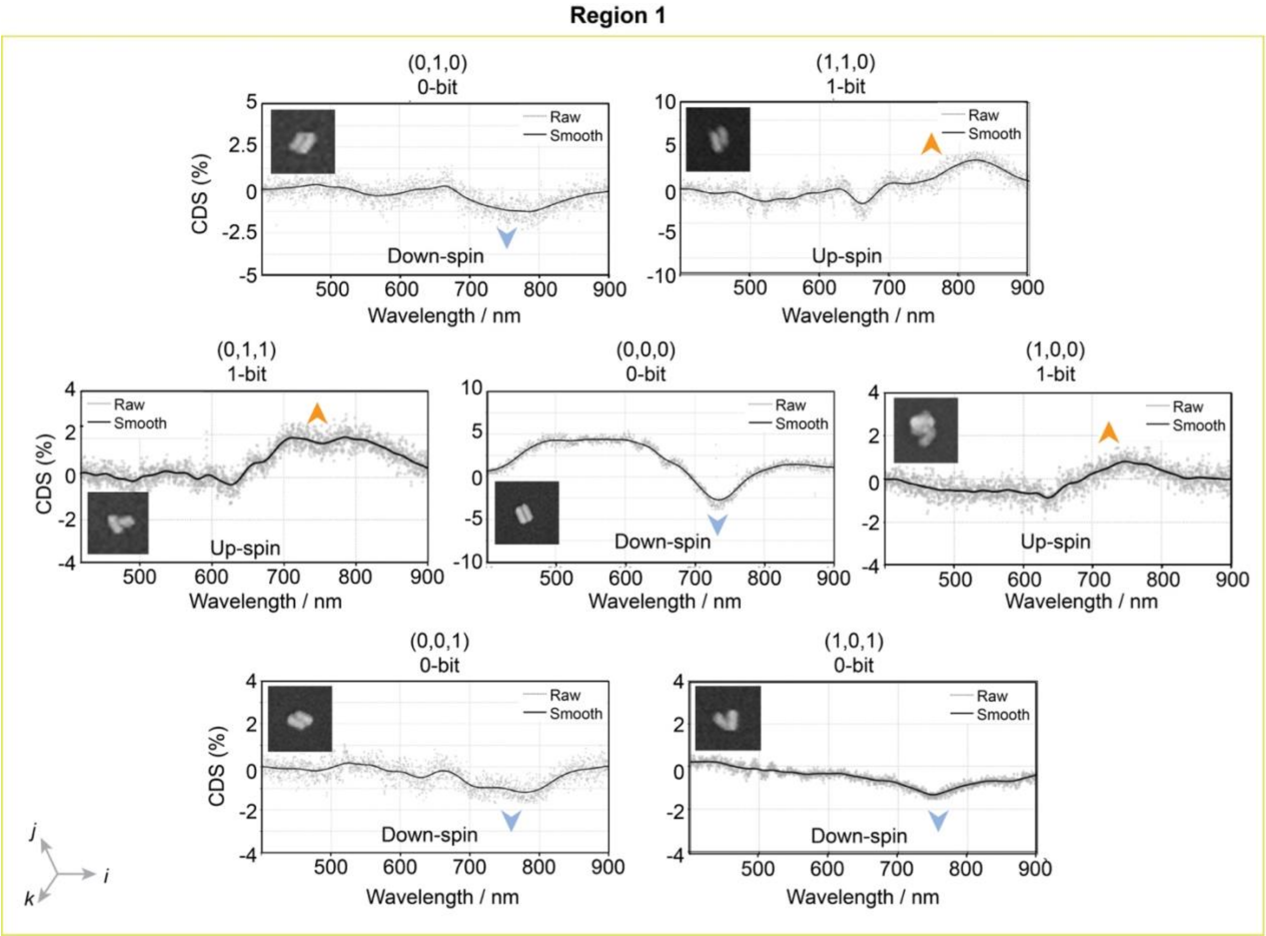


**Figure S14. Additional CD spectra collected from nanoarray region 1.** Up-spin = 1-bit and down-spin = 0-bit.

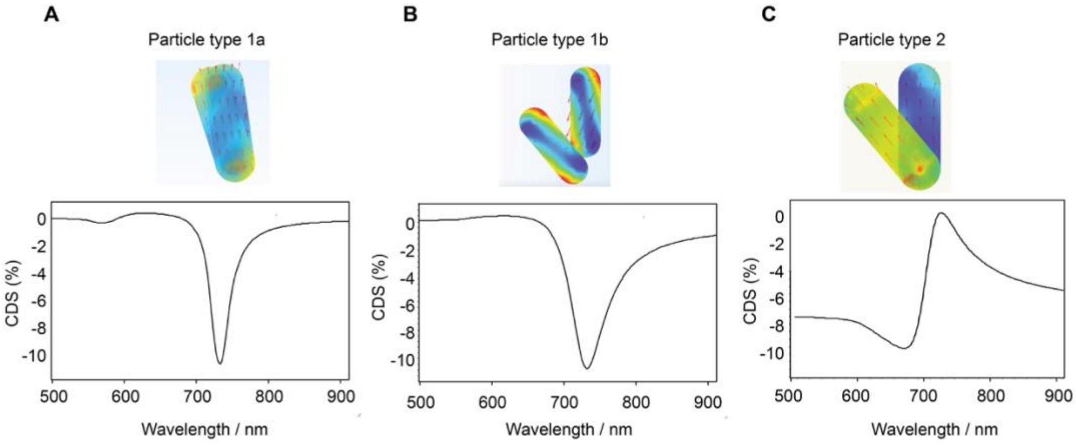


**Figure S15. Electromagnetic Waves, Frequency Domain (EWFD) simulations.** (A) Almost parallel geometry shows a single peak dip. (B) V-shape arrangement with weaker dipole interaction also shows a single peak. (C) Almost perfectly aligned AuNRs with angle more than 20º shows a bisignate feature.

**Section 2. DNA sequences**

**Table S1.** Various buffer recipes used in this work.

| Solution | Composition | pH | Role |
|---|---|---|---|
| Hybridization buffer | 12.5 mM MgCl2, 40 mM Tris-HCl, EDTA | 8.4 | Formation of triangle origami by hybridization of scaffold and staples |
| Binding buffer | 1 mM NiCl2, 40 mM Tris-HCl | 6.5 | Improves placement of origami on binding spots avoiding hydrolysis of HMDS |
| Buffer 1 | 40 mM MgCl2, 40 mM Tris-HCl | 8.4 | Builds interaction of triangle origami with hydrophilic binding spots |
| Buffer 2 | 40 mM MgCl2, 40 mM Tris-HCl, 0.07% tween 20 | 8.4 | Hydrates the surface and repositions origami in binding spots, also removes origami bound on hydrophobic HMDS |
| Buffer 3 | 35 mM MgCl2, 40 mM Tris-HCl | 8.9 | Hydrolysis HMDS and removes non-specifically bound origami |
| Silicification buffer | 12.5 mM MgCl2, 40 mM Tris-HCl, 2mM EDTA | 8.0 | Provides both hybridization buffer conditions to origami while maintaining its intactness |

**Table S2.** DNA strand sequences used for attaching gold nanorods and DNA-PAINT or photobleaching experiments.

| Experiment | Description | Sequence |
|---|---|---|
| Chiral metamaterial | Top-handle | AAA AAA AAA AAA AAA/3ThioMC3-D/ |
| | Bottom-handle | TTT ATT TAT TAT TTT AAA 3ThioMC3-D/ |
| DNA-PAINT and Photobleaching (12 handles across three edges) | Imager-PAINT | AGG AGG A /5AmMC6/ |
| | Imager-bleaching | AGG AGG AGG AGG AGG AGG A/3AmMO/cy3b |
| | strand_51-docking | TTGCAAAATATAGTCAGAAGCAAAACCTTTAA**TCCTCCTCCTCCTCCTCCT** |
| | strand_54-docking | TGCGGAATGCTTTAAACAGTTCAGTATAATGC**TCCTCCTCCTCCTCCTCCT** |
| | strand_58-docking | TTGACCCCTCCATTAAACGGGTAACAGCTTGC**TCCTCCTCCTCCTCCTCCT** |
| | strand_61-docking | TAAATTGTTCGGAACGAGGGTAGCCAACCATC**TCCTCCTCCTCCTCCTCCT** |
| | strand_125-docking | ATAGCTATAGGTGGCAACATATAAAAATATTG**TCCTCCTCCTCCTCCTCCT** |
| | strand_128-docking | GGAAACCGAACTGGCATGATTAAGAATTAGAG**TCCTCCTCCTCCTCCTCCT** |
| | strand_132-docking | AACAATAGATAAAGTACCGACAAATAAATTTA**TCCTCCTCCTCCTCCTCCT** |
| | strand_135-docking | CAATCAATCCATATTTAACAACGCGAATCATA**TCCTCCTCCTCCTCCTCCT** |
| | strand_195-docking | GTGAGGCCCTCATGGAAATACCTACGTAAGAA**TCCTCCTCCTCCTCCTCCT** |
| | strand_198-docking | GATTAGTAACTATCGGCCTTGCTGAGCCCTAA**TCCTCCTCCTCCTCCTCCT** |
| | strand_202-docking | ACCTGTCGAGCATAAAGTGTAAAGATTACGCC**TCCTCCTCCTCCTCCTCCT** |
| | strand_205-docking | GCGCCAGGGTCATAGCTGTTTCCTTCCCAGTC**TCCTCCTCCTCCTCCTCCT** |
| | strand_35-docking | **/5Biosg/tttt**TTAAATATTCCAAAAGGAGCCTTTTCTCCAAA |
| | strand_44-docking | **/5Biosg/tttt**GAGGTCATTCTTTACCCTGACTATGAAGTTTTGCCAGAGGTATACCAG |
| | strand_45-docking | **/5Biosg/tttt**TTAAACAGTTTTCATGAGGAAGTTCAGCGATTATACCAAGGCTCATTC |
| | strand_110-docking | **/5Biosg/tttt**TAGCACCAAAATATATTTTAGTTAGCGAGAAA |
| | strand_118-docking | **/5Biosg/tttt**AAAGGTGAGAAAATACATACATAACTTACCGAAGCCCTTTTTATCCCA |
| | strand_119-docking | **/5Biosg/tttt**ACCGTGTGCCAGTAATAAGAGAATATAAGTCCTGAACAAGTTGAAGCC |
| | strand_181-docking | **/5Biosg/tttt**ATACCGAATGGGAAGGGCGATCGGCAGGCTGC |
| | strand_189-docking | **/5Biosg/tttt**TATTTTTGTGCAACAGGAAAAACGACCGAGTAAAAGAGTCCGGGCGCT |
| | strand_190-docking | **/5Biosg/tttt**ATGTGCTGCAACATACGAGCCGGATGCCAGCTGCATTAATGGAACAAG |

**Table S3.** DNA sequences used to make triangular origami.

| Start | End | Sequence | Length | Color |
|---|---|---|---|---|
| 20[255] | 22[256] | TGGAGCAATAAGCAAATATTTAAACATTTTTT | 32 | #55aa00 |
| 3[112] | 2[109] | ATCAATTCAGGATAAAAAT | 19 | #000000 |
| 10[239] | 9[223] | CGTCAGATGAATATACAGTAACAGTCGCCTGA | 32 | #aa5500 |
| 4[287] | 6[288] | CAGTTTCAGCGCCGACAATGACAAAACGGCTA | 32 | #000000 |
| 19[304] | 3[90] | ATCGTAACCGTGCATCTGCATATTTTCATTTGG | 33 | #000000 |
| 9[160] | 11[159] | TACCGTTCGCCGCCGCCAGCATTGCCTCCCTC | 32 | #aa5500 |
| 4[159] | 6[160] | CAGTTGATGAGCTTAATTGCTGAAAAAACGAG | 32 | #000000 |
| 16[63] | 18[64] | CAATCAATAACCCACAAGAATTGACAGAGAGA | 32 | #000000 |
| 6[223] | 8[224] | ATGCCACTAAGAATACACTAAAACGAAACACC | 32 | #000000 |
| 6[127] | 8[128] | CTGACTATGAAGTTTTGCCAGAGGTATACCAG | 32 | #000000 |
| 14[239] | 12[240] | AAATCCAATAAATCGTCGCTATTATTAACAAT | 32 | #aa5500 |
| 23[304] | 21[303] | GCGCCAGGGTCATAGCTGTTTCCTTCCCAGTC | 32 | #000000 |
| 0[255] | 2[256] | ACATGAAAGTATAGCCCGGAATAGCAGAACCG | 32 | #000000 |
| 10[271] | 9[255] | AATAAAGAAATTGCGTAGATTTTCATCGCGCA | 32 | #aa5500 |
| 21[304] | 19[303] | ACGACGTTGACGACGACAGTATCGATGGGCGC | 32 | #000000 |
| 8[287] | 7[271] | TACCCAAATCAACGTAACAAAGCTCGCGAAAC | 32 | #000000 |
| 1[280] | 9[119] | AGGAGGTTTAGAAACAAATAAA | 22 | #000000 |
| 13[288] | 15[287] | AAACACCGCAACATGTAATTTAGGCCTAATTT | 32 | #aa5500 |
| 26[95] | 28[96] | CGCTCAATTCCTGAGAAGTGTTTTACGCTGCG | 32 | #55aa00 |
| 22[191] | 19[207] | AGGATTTAAGCCGTCAATAGATAATCATCAAC | 32 | #55aa00 |
| 4[191] | 5[207] | AGTACGGTAACATGTTTTAAATATTCCAAAAG | 32 | #000000 |
| 15[288] | 18[272] | ACGAGCATGTTTTAGCGAACCTCCCGACTTGC | 32 | #000000 |
| 28[95] | 23[79] | CGTAACCACCACACCCGCCGCGCTGAACGGTA | 32 | #000000 |
| 3[240] | 1[239] | ACGCCTGTAGAGCCACCACCCTCAGAGAGGGT | 32 | #000000 |
| 6[319] | 8[320] | AGACAGCAGTCGAAATCCGCGACCCCTTCATC | 32 | #000000 |
| 0[191] | 1[207] | TATGATATCGGAGACAGTCAAATCTTTTGCTC | 32 | #000000 |
| 19[240] | 17[239] | TCTCCGTGGGAACGCCATCAAAAAAAACAGGA | 32 | #55aa00 |
| 18[63] | 15[47] | ATAACATAAAAACAGGGAAGCGCACTAATATC | 32 | #000000 |
| 7[304] | 5[303] | TAAATTGTTCGGAACGAGGGTAGCCAACCATC | 32 | #000000 |
| 2[255] | 4[256] | CCACCCTCAGCATTCCACAGACAGAAGGAACA | 32 | #000000 |
| 5[304] | 3[303] | GCCCACGCGGGATTTTGCTAAACAGTCGTCTT | 32 | #000000 |
| 5[112] | 3[111] | TTGCTCCTACCATTAGATACATTTAAGGTGGC | 32 | #000000 |
| 1[240] | 0[224] | TGATATAAGTATTAAGAGGCTGAGACTCCTCA | 32 | #000000 |
| 13[160] | 15[159] | CATTTGGGACTCCTTATTACGCAGGATAGCCG | 32 | #aa5500 |
| 17[240] | 20[224] | AGATTGTAACAAGAGAATCGATGAACGGTAAT | 32 | #55aa00 |
| 6[287] | 8[288] | CAGAGGCTACGGAGATTTGTATCAATATTCAT | 32 | #000000 |
| 15[256] | 18[240] | TGAACAAGTTGAAGCCTTAAATCAAGATTAGT | 32 | #000000 |
| 21[240] | 19[239] | AGCTGGCGGCCGGAAACCAGGCAACGTCGGAT | 32 | #55aa00 |

| | | | | |
|---|---|---|---|---|
| 8[319] | 7[303] | AAGAGTAATCTTGACAAGAACCGGTCGCCTGA | 32 | #000000 |
| 12[207] | 11[191] | TACATAAACCCTCAGAGCCGCCACACCCTCAG | 32 | #aa5500 |
| 11[192] | 12[208] | AGCCACCATCAATATATGTGAGTGGGAAACAG | 32 | #aa5500 |
| 9[192] | 10[208] | GATGATACAGAAACAATAACGGATTACCTTTT | 32 | #aa5500 |
| 8[127] | 7[111] | TCAGGACGTTGGGAAGAAAAATCTAGAGGCTT | 32 | #000000 |
| 26[63] | 28[64] | GGCAGATTAAGGGATTTTAGACAGTAATGCGC | 32 | #000000 |
| 12[298] | 11[287] | TTTTTTTTGAAGATGATGAGCGATAGC | 27 | #000000 |
| 23[80] | 21[79] | CGCCAGAACGTCTGAAATGGATTACCAACAGA | 32 | #000000 |
| 11[308] | 19[90] | GAGTCAATAGTGAATTATCAAACCCTCAA | 29 | #000000 |
| 24[255] | 26[256] | CTTCTGGTAAAGGGGGATGTGCTGCAACATAC | 32 | #55aa00 |
| 18[79] | 16[80] | AGCCTTTAGTTAAGCCCAATAATACCACGGAA | 32 | #000000 |
| 13[256] | 15[255] | ACCGTGTGCCAGTAATAAGAGAATATAAGTCC | 32 | #aa5500 |
| 24[327] | 26[320] | TTTTTCAGTTTGAGGGGTAAAACGACGGCCAGCGAATTCG | 40 | #000000 |
| 28[351] | 23[335] | ATCCTGTTTGATGGTGGTTCCGAACCAGTGAG | 32 | #000000 |
| 26[127] | 28[128] | GAAAAACGACCGAGTAAAAGAGTCCGGGCGCT | 32 | #55aa00 |
| 6[159] | 8[160] | AATGACCATTTAGACTGGATAGCGATTACCTT | 32 | #000000 |
| 15[96] | 18[80] | AAACAATGTTTAACGTCAAAAATGAAAATAGC | 32 | #000000 |
| 4[95] | 6[96] | GTCAATAAGTCAGGATTAGAGAGTGCGGATTG | 32 | #000000 |
| 2[159] | 4[160] | CCTGTAATTCATACAGGCAAGGCACCATATAA | 32 | #000000 |
| 21[59] | 26[64] | AGGGACATTCTGGTTTACATT | 21 | #000000 |
| 17[176] | 20[160] | GATTATCAGTTTGGATTATACTTCTGAATAAT | 32 | #55aa00 |
| 23[240] | 21[239] | ACCTGTCGAGCATAAAGTGTAAAGATTACGCC | 32 | #000000 |
| 11[224] | 13[223] | TGCTTCTGTCGCAAGACAAAGAACATTTCATC | 32 | #aa5500 |
| 16[79] | 14[77] | TAAGTTTAGGCGACATTCAACCGACGGCATTTTCG | 35 | #000000 |
| 26[255] | 28[256] | GAGCCGGATGCCAGCTGCATTAATGGAACAAG | 32 | #55aa00 |
| 7[80] | 5[79] | ACGACGATAGATTAAGAGGAAGCCAAGCAAAC | 32 | #000000 |
| 15[224] | 18[192] | AACGCGCCTGCACCCAGCTACAATTTTATCCTGAATCTTAC CAACGCT | 48 | #000000 |
| 7[336] | 5[335] | GTTACTTAGGGATCGTCACCCTCAGTCGCTGA | 32 | #000000 |
| 16[239] | 14[240] | CCGACAAATAAATTTAATGGTTTGGCTGATGC | 32 | #aa5500 |
| 18[111] | 16[112] | ATTTTTTGAAATAGCAATAGCTATAGGTGGCA | 32 | #000000 |
| 14[175] | 12[176] | TCACCAATCACCCTCAGAACCGCCCAGAACCA | 32 | #aa5500 |
| 22[223] | 24[224] | GTCTGGCCTGAGCGAGTAACAACCAGCGCCAT | 32 | #55aa00 |
| 28[319] | 23[303] | AATCCCTTATAAATCAAAAGAATACGTATTGG | 32 | #000000 |
| 1[176] | 0[160] | AGAAAGGCTCAACCGTTCTAGCTGATAAATTA | 32 | #000000 |
| 2[295] | 4[288] | TTTTTTACCGCCACCCAACGATCTAAAGTTTTACTTTCAA | 40 | #000000 |
| 7[240] | 5[239] | TTGACCCCTCCATTAAACGGGTAACAGCTTGC | 32 | #000000 |
| 6[255] | 8[256] | AGGAAGTTCAGCGATTATACCAAGGCTCATTC | 32 | #000000 |
| 14[271] | 12[272] | TGGGTTATAATCCTTGAAAACATAAACAAACA | 32 | #aa5500 |
| 4[223] | 6[224] | GTTGAAAAAATTGTATCGGTTTATAATACGTA | 32 | #000000 |
| 15[27] | 18[13] | GGGTAATTGAGCGTTAGACGGGAGAATTAACTGAACACCC | 40 | #000000 |
| 20[223] | 22[224] | CGTAAAACATCAGAAAAGCCCCAATAATTCGC | 32 | #55aa00 |

| | | | | |
|---|---|---|---|---|
| 5[336] | 13[58] | GGCTTGCAGGGAGTTAAAGATGGTTTACCAGCG | 33 | #000000 |
| 5[240] | 3[239] | TTTCGAGGAATTGCGAATAATAATAAACTACA | 32 | #000000 |
| 5[80] | 4[77] | TCCAACAGCCTGTTTAGCT | 19 | #000000 |
| 8[223] | 8[192] | AGAACGAGTAGTAAATTGGGCTTGAGATGGTT | 32 | #000000 |
| 26[287] | 28[288] | TTGTTATCGGGGAGAGGCGGTTTGGCCCGAGA | 32 | #000000 |
| 24[191] | 21[207] | CAGCAGAACATTAAAAATACCGAATGGGAAGG | 32 | #55aa00 |
| 11[96] | 13[95] | CGTTTGCCTGTAGCGCGTTTTCATTTGAGGGA | 32 | #000000 |
| 10[175] | 9[159] | AACGGGGTCAGTGCCTTGAGTAACTCTGAATT | 32 | #aa5500 |
| 7[368] | 15[26] | ATCATAAGGGAACCGAACTTGAACAAAGTCAGA | 33 | #000000 |
| 21[144] | 19[143] | TTAGTCTTACCGCCTGCAACAGTGGAAGGTTA | 32 | #55aa00 |
| 28[386] | 23[367] | AAGCGGTCCACGCTGGTTTGCCCCAGCACCGCCTG | 35 | #000000 |
| 15[128] | 18[112] | AGCCCTTTTTATCCCAATCCAAATAAGAAACG | 32 | #000000 |
| 10[143] | 9[127] | TATAAACAGTTAATGCCCCCTGCCTCCTCATT | 32 | #aa5500 |
| 15[192] | 16[208] | CAATAATAAAACAACATGTTCAGCTCCAGACG | 32 | #000000 |
| 17[280] | 1[119] | TTTGTTAAAATTTTTAGAACCC | 22 | #000000 |
| 16[207] | 15[191] | ACGACAATACGGAATACCCAAAAGAGGAAACG | 32 | #aa5500 |
| 3[144] | 1[143] | CCAATAAAACTTTTGCGGGAGAAGCAATGCCT | 32 | #000000 |
| 11[288] | 13[287] | TTAGATTAACCTTTTTAACCTCCGATAAGAAT | 32 | #aa5500 |
| 16[271] | 14[272] | TTTTCGAGATAAATAAGGCGTTAAGCTTAGGT | 32 | #aa5500 |
| 13[224] | 15[223] | TTCTGACCAGGTAAAGTAATTCTGTAATGCAG | 32 | #aa5500 |
| 2[191] | 3[207] | AGCTAAATAAGCAATAAAGCCTCATGTACCGT | 32 | #000000 |
| 13[59] | 16[64] | CCAAAGACAAAAGTTTTGTCA | 21 | #000000 |
| 26[359] | 28[352] | TTTTTCTCTAGAGGATCAGCTGATTGCCCTTCAGGCGAAA | 40 | #000000 |
| 9[224] | 11[223] | TTGCTTTGAATTACCTTTTTTAATAATAACCT | 32 | #aa5500 |
| 8[159] | 7[143] | ATGCGATTTTAAGAACTGGCTCATGGGTAATA | 32 | #000000 |
| 5[144] | 3[143] | ATGGCTTATCCCAATTCTGCGAACATTAACAT | 32 | #000000 |
| 28[207] | 23[175] | GTCTATCATTTAGAGCTTGACGGGGAAAGCCGGCGAACGT ACTTCTTT | 48 | #000000 |
| 5[272] | 3[271] | CGATAGTTGCGGAGTGAGAATAGACCCTCATA | 32 | #000000 |
| 13[192] | 14[208] | TAGCACCAAAATATATTTTAGTTAGCGAGAAA | 32 | #aa5500 |
| 22[295] | 24[288] | TTTTTTCGCATTAAATGGTCACGTTGGTGTAGGCCTCAGG | 40 | #000000 |
| 0[159] | 2[160] | ATGCCGGATGTAGGTAAAGATTCAATTATGAC | 32 | #000000 |
| 7[144] | 5[143] | GTAAAATGTAAATCAAAAATCAGGTTTTGCGG | 32 | #000000 |
| 7[48] | 6[45] | AGAGCAACTCGCGTTTTAA | 19 | #000000 |
| 18[239] | 16[240] | TGCTATTTTGTTTATCAACAATAGATAAAGTA | 32 | #000000 |
| 22[255] | 24[256] | AACCAATAGGAACAAACGGCGGATCGGCACCG | 32 | #55aa00 |
| 24[223] | 26[224] | TCGCCATTTGCGGGCCTCTTCGCTCCTGGGGT | 32 | #55aa00 |
| 10[207] | 9[191] | ACATCGGGAGGAGTGTACTGGTAATGGCTTTT | 32 | #aa5500 |
| 7[272] | 5[271] | AAAGTACATTGAGGACTAAAGACTCTTGATAC | 32 | #000000 |
| 3[272] | 1[271] | GTTAGCGTTCAGAACCGCCACCCTGTGTATCA | 32 | #000000 |
| 14[143] | 12[144] | ATCAGTAGGCCACCACCGGAACCGACAGGAGG | 32 | #aa5500 |
| 2[223] | 4[224] | GATAGCAAGTTTCGTCACCAGTACTTTTTCAC | 32 | #000000 |

| | | | | |
|---|---|---|---|---|
| 12[271] | 10[272] | TCAAGAAATTATTCATTTCAATTAAAAACAGA | 32 | #aa5500 |
| 7[208] | 6[192] | AGAGGCAAACGAAGGCACCAACCTTGAATCCC | 32 | #000000 |
| 14[207] | 13[191] | ACTTTTTCTTACCATTAGCAAGGCATCACCAG | 32 | #aa5500 |
| 1[208] | 0[192] | AGTACCAGATTAGGATTAGCGGGGACCATCAA | 32 | #000000 |
| 24[287] | 26[288] | AAGATCGCGGTAACGCCAGGGTTTGTGTGAAA | 32 | #000000 |
| 19[91] | 24[96] | TCAATATCTGGTCTAAAGCAT | 21 | #000000 |
| 12[143] | 10[144] | TTGAGGCAAATGGAAAGCGCAGTCAGTGCCCG | 32 | #aa5500 |
| 15[368] | 23[26] | CATCGAGAACAAGCAAGCCTCCTCGTTAGAATC | 33 | #000000 |
| 7[176] | 5[175] | TGCGGAATGCTTTAAACAGTTCAGTATAATGC | 32 | #000000 |
| 12[239] | 10[240] | TTCATTTGAATACCAAGTTACAAAAGGTTTAA | 32 | #aa5500 |
| 17[272] | 20[256] | GTTAATATCTATCAGGTCATTGCCTGAGAGTC | 32 | #55aa00 |
| 13[336] | 21[58] | CGTTATACAAATTCTTACCCGACCAGTAATAAA | 33 | #000000 |
| 21[112] | 19[111] | TACGTGGCCAGCAAATGAAAAATCAGTTGGCA | 32 | #55aa00 |
| 6[191] | 7[207] | CCTCAAATCGTCATAAATATTCATAAAACGAA | 32 | #000000 |
| 1[272] | 0[256] | CCGTACTCTATTTCGGAACCTATTATTCTGAA | 32 | #000000 |
| 4[326] | 6[320] | TTTTATTTTCTGTATATAACCGATATATTCGGCAGCGAA | 39 | #000000 |
| 17[208] | 20[192] | GGTTGATATAGCATGTCAATCATACAATATAA | 32 | #55aa00 |
| 16[111] | 14[112] | ACATATAAAAATATTGACGGAAATGCCTTTAG | 32 | #000000 |
| 9[128] | 11[127] | AAAGCCAGGGTCAGACGATTGGCCAATCACCG | 32 | #aa5500 |
| 15[160] | 18[144] | AACAAAGTTTTGCCAGTTACAAAATAAACAGC | 32 | #000000 |
| 23[272] | 21[271] | CAACGCGCCGCTCACAATTCCACACAAGGCGA | 32 | #000000 |
| 21[208] | 24[192] | GCGATCGGCAGGCTGCGCAACTGTCGAACCAC | 32 | #55aa00 |
| 6[359] | 8[352] | TTTTTGCCGCTTTTGCGCCGGAACGAGGCGCACGGTGTAC | 40 | #000000 |
| 11[256] | 13[255] | TCCCTTAGATAACTATATGTAAATAAATACCG | 32 | #aa5500 |
| 21[272] | 19[271] | TTAAGTTGACTCCAGCCAGCTTTCTGACCGTA | 32 | #55aa00 |
| 18[386] | 15[367] | GTTTTTATTTTCATCGTAGGAATCATTACCGCACT | 35 | #000000 |
| 16[143] | 14[144] | CAAACGTAATTATCACCGTCACCGGCACCGTA | 32 | #aa5500 |
| 28[63] | 23[47] | CGCTACAGGGCGCGTACTATGGTTAACAGGAG | 32 | #000000 |
| 22[127] | 24[128] | GTTATTAAGTTGAAAGGAATTGAGCCACGCTG | 32 | #55aa00 |
| 15[336] | 13[335] | CCAAGAACAACGCTCAACAGTAGGATCATATG | 32 | #000000 |
| 24[159] | 26[160] | GTATTAACTAATGCGCGAACTGATGTAATATC | 32 | #55aa00 |
| 5[59] | 6[64] | CGAACCAGACCGGCGAAAGAC | 21 | #000000 |
| 23[368] | 7[26] | GCCCTGAGAGAGTTTTTTATACATAACGCCAA | 33 | #000000 |
| 8[191] | 7[175] | TAATTTCAACTTTAATCATTGTGATCCAATAC | 32 | #000000 |
| 5[176] | 3[175] | TGTAGCTCGTCTGGAAGTTTCATTAAGAATTA | 32 | #000000 |
| 16[319] | 18[320] | GAGAATCGAATCGGCTGTCTTTCCAAGGCTTA | 32 | #aa5500 |
| 23[112] | 21[111] | GTGAGGCCCTCATGGAAATACCTACGTAAGAA | 32 | #000000 |
| 4[127] | 6[128] | TTAGTTTGTTTGATAAGAGGTCATTCTTTACC | 32 | #000000 |
| 4[255] | 6[256] | ACTAAAGGTGAATTTCTTAAACAGTTTTCATG | 32 | #000000 |
| 18[319] | 16[304] | TCCGGTATTCTAAGAACGCGAGGCGTAGAAACCAATCAAT CCATATTT | 48 | #000000 |
| 19[144] | 17[143] | TCTAAAATTCGTATTAAATCCTTTAAGAAACC | 32 | #55aa00 |

| | | | | |
|---|---|---|---|---|
| 3[91] | 4[96] | GGCGCGAGCTGAACGCAAATG | 21 | #000000 |
| 19[176] | 17[175] | TAGATTAGGAAGTATTAGACTTTAATATTCCT | 32 | #55aa00 |
| 19[112] | 22[109] | AATCAACATTTTAAAAGTT | 19 | #55aa00 |
| 19[208] | 22[192] | ATTAAATGTTCCTGTAGCCAGCTTTACATTTG | 32 | #55aa00 |
| 3[208] | 2[192] | AACACTGAGCCCAATAGGAACCCAGAGCATAA | 32 | #000000 |
| 0[125] | 2[128] | CAAAGGTCATATATTTTAAATGCCTTTATT | 30 | #000000 |
| 16[303] | 14[304] | AACAACGCGAATCATAATTACTAGATAGGTCT | 32 | #aa5500 |
| 19[272] | 17[271] | ATGGGATATTTTGTTAAATCAGCTTTGTAAAC | 32 | #55aa00 |
| 0[223] | 2[224] | AGAGAAGGGCGGATAAGTGCCGTCTTTTCAGG | 32 | #000000 |
| 14[327] | 16[320] | TTTTTTTATCAAAATCAAAAAGCCTGTTTAGTGCTTAATT | 40 | #000000 |
| 14[303] | 11[307] | GAGAGACTAGACGCTGAGAA | 20 | #aa5500 |
| 24[95] | 26[96] | CACCTTGCCCTTCTGACCTGAAAGCATTTGA | 32 | #55aa00 |
| 20[159] | 22[160] | GGAAGGGTGAGCGGAATTATCATCCAAACAAT | 32 | #55aa00 |
| 18[351] | 15[335] | CAATAGCAAGCAAATCAGATATAGTTATCATT | 32 | #000000 |
| 18[191] | 16[176] | AACGAGCGTCTTTCCAGAGCCTAATACCAGAAGGAAACCG AACTGGCA | 48 | #000000 |
| 24[127] | 26[128] | AGAGCCAGACAGACAATATTTTTGTGCAACAG | 32 | #55aa00 |
| 16[175] | 14[176] | TGATTAAGAATTAGAGCCAGCAAACGGAAACG | 32 | #aa5500 |
| 8[386] | 7[367] | GACCAACTTTGAAAGAGGACAGATGAAGACGGTCA | 35 | #000000 |
| 16[359] | 18[352] | TTTTTAGTATAAAGCCGGGTATTAAACCAAGTACCGCGCC | 40 | #000000 |
| 7[27] | 8[13] | AAGGAATTACGAGTAGGAATACCACATTCAACTAATGCAG | 40 | #000000 |
| 23[48] | 26[45] | GCCGATTACACCAGTCACA | 19 | #000000 |
| 28[159] | 23[143] | AGGAAGGGAAGAAAGCGAAAGGAGTGTCCATC | 32 | #000000 |
| 26[223] | 28[208] | GCCTAATGCACTGCCCGCTTTCCATCCAACGTCAAAGGGCG AAAAACC | 48 | #55aa00 |
| 21[336] | 5[58] | TTGCATGCCTGCAGGTCTTTTCGAGCTTCAAAG | 33 | #000000 |
| 18[271] | 16[272] | GGGAGGTTAAAAATAATATCCCATCAGAGGCA | 32 | #000000 |
| 7[112] | 5[111] | TTGCAAAATATAGTCAGAAGCAAAACCTTTAA | 32 | #000000 |
| 6[63] | 8[64] | TTCAAATAACTATCATAACCCTCGATTACAGG | 32 | #000000 |
| 20[127] | 22[128] | TTGCACGTATCATTTTGCGGAACAGCCCGAAC | 32 | #55aa00 |
| 22[159] | 24[160] | TCGACAACATCTTTAGGAGCACTAGGCGGTCA | 32 | #55aa00 |
| 28[255] | 23[239] | AGTCCACTATTAAAGAACGTGGACGTCGGGAA | 32 | #000000 |
| 17[144] | 20[128] | ACCAGAAGTAGAACCTACCATATCAAAATTAT | 32 | #55aa00 |
| 14[111] | 12[109] | CGTCAGACATCTTTTCATAATCAATTGATATTCAC | 35 | #000000 |
| 13[96] | 15[95] | GGGAAGGTAAGAAACGCAAAGACAAGAGCAAG | 32 | #000000 |
| 11[160] | 13[159] | AGAGCCGCGAAACCATCGATAGCAACTTGAGC | 32 | #aa5500 |
| 23[208] | 26[192] | GTTGCGCTAGTGAGCTAACTCACAGAGTAGAA | 32 | #000000 |
| 26[159] | 28[160] | CAGAACAATAACCGTTGTAGCAATGGCGAGAA | 32 | #55aa00 |
| 21[80] | 24[77] | GATAGAACTGAACCTCAAA | 19 | #55aa00 |
| 3[176] | 1[175] | GCAAAATTCGGTTGTACCAAAAACAAAGGGTG | 32 | #000000 |
| 26[319] | 28[320] | TAATCATGGTGGTTTTTCTTTTCAATCGGCAA | 32 | #000000 |
| 9[256] | 11[255] | GAGGCGAAACAAAATTAATTACATATTAATTT | 32 | #aa5500 |
| 21[176] | 19[175] | AACATCGCGATAAAACAGAGGTGAACAACTAA | 32 | #55aa00 |

| | | | | |
|---|---|---|---|---|
| 8[351] | 7[335] | AGACCAGGCGCATAGGCTGGCTGATGCTCCAT | 32 | #000000 |
| 18[143] | 16[144] | CATATTATTTAAGAAAAGTAAGCATATGTTAG | 32 | #000000 |
| 20[191] | 17[207] | TCCTGATTGATGATGGCAATTCATTGTACCCC | 32 | #55aa00 |
| 9[280] | 17[119] | CCTGAGCAAAATGAGTAACATT | 22 | #000000 |
| 8[255] | 7[239] | AGTGAATAAGGCTTGCCCTGACGAACTCATCT | 32 | #000000 |
| 6[95] | 8[96] | CATCAAAAAAAAACCAAAATAGCGACGTTAAT | 32 | #000000 |
| 3[304] | 11[95] | TCCAGACGTTAGTAAATGAGTCATAGCCCCCTTATTAG | 38 | #000000 |
| 23[336] | 21[335] | ACGGGCAACCCCGGGTACCGAGCTTGCCAAGC | 32 | #000000 |
| 5[208] | 4[192] | GAGCCTTTTCTCCAAAAAAAAGGCGCAACTAA | 32 | #000000 |
| 2[127] | 4[128] | TCAACGCATACTAATAGTAGTAGCGAGTAGAT | 32 | #000000 |
| 28[287] | 23[271] | TAGGGTTGAGTGTTGTTCCAGTTTGAATCGGC | 32 | #000000 |
| 23[27] | 28[13] | AGAGCGGGAGCTAGCTTTGACGAGCACGTATAACGTGCTT | 40 | #000000 |
| 1[144] | 0[126] | GAGTAATGGAGGGTAGCTATTTTTGAGAGATCTA | 34 | #000000 |
| 8[95] | 7[79] | AAAACGAACTAACGGAACAACATTTTTACCAG | 32 | #000000 |
| 13[128] | 15[127] | AAAGGTGAGAAAATACATACATAACTTACCGA | 32 | #aa5500 |
| 28[127] | 23[111] | AGGGCGCTGGCAAGTGTAGCGGTCTATAATCA | 32 | #000000 |
| 23[176] | 21[175] | GATTAGTAACTATCGGCCTTGCTGAGCCCTAA | 32 | #000000 |
| 12[175] | 10[176] | CCACCAGACAGTAAGCGTCATACATAAGTTTT | 32 | #aa5500 |
| 11[128] | 13[127] | GAACCAGACGACAGAATCAAGTTTTATTCATT | 32 | #aa5500 |
| 8[63] | 7[47] | TAGAAAGATTCATCAGTTGAGATTGCATAGTA | 32 | #000000 |
| 26[191] | 23[207] | GAACTCAAATAACATCACTTGCCTTTAATTGC | 32 | #55aa00 |
| 23[144] | 21[143] | ACGCAAATTATTACCGCCAGCCATAATGGCTA | 32 | #000000 |
| 15[48] | 16[45] | AGAGAGATAGAAAATTCAT | 19 | #000000 |
